# Defining short linear motif binding determinants by phage-based multiplexed deep mutational scanning

**DOI:** 10.1101/2024.08.06.606761

**Authors:** Caroline Benz, Lars Maasen, Leandro Simonetti, Filip Mihalic, Richard Lindqvist, Ifigenia Tsitsa, Per Jemth, Anna K. Överby, Norman E. Davey, Ylva Ivarsson

## Abstract

Deep mutational scanning (DMS) has emerged as a powerful approach for evaluating the effects of mutations on binding or function. Here, we developed a multiplexed DMS by phage display protocol to define the binding determinants of short linear motifs (SLiMs) binding to peptide binding domains. We first designed a benchmarking DMS library to evaluate the performance of the approach on well-known ligands for eleven different peptide binding domains, including the talin-1 PTB domain. Systematic benchmarking against a gold-standard set of motifs from the eukaryotic linear motif (ELM) database confirmed that the DMS by phage analysis correctly identifies known motif binding determinants. The DMS analysis further defined a non-canonical PTB binding motif, with a putative extended conformation. A second DMS library was designed aiming to provide information on the binding determinants for 19 SLiM-based interactions between human and SARS-CoV-2 proteins. The analysis confirmed the affinity determining residues of viral peptides binding to host proteins, and refined the consensus motifs in human peptides binding to five domains from SARS-CoV-2 proteins, including the non-structural protein (NSP) 9. The DMS analysis further pinpointed mutations that increased the affinity of ligands for NSP3 and NSP9. An affinity improved cell-permeable NSP9-binding peptide was found to exert stronger antiviral effects as compared to the initial wild-type peptide. Our study demonstrates that DMS by phage display can efficiently be multiplexed and applied to refine binding determinants, and shows how DMS by phage display can guide peptide-engineering efforts.

## Introduction

Short linear motifs (SLiMs) are compact protein-protein interaction modules typically found in the intrinsically disordered regions (IDRs) of the proteome (Tompa, Davey et al. 2014). SLiM-based interactions play a crucial role in several important cellular processes such as signal transduction, enzyme recruitment for activation of the protein, and protein localization. A general picture of SLiM-based interactions has emerged where a limited set of 3-4 key residues serve as the main specificity and affinity determinants, and where motif-flanking regions may contribute and modulate binding (Holehouse and Kragelund 2024, Kumar, Michael et al. 2024, Mihalic, Arcila et al. 2024). Disease-associated mutations in the IDRs have been found to both break and make SLiM-based interactions (Meszaros, Kumar et al. 2017, Kliche, Simonetti et al. 2024, Rrustemi, Meyer et al. 2024). Furthermore, viruses also exploit SLiM-based interactions to outcompete the endogenous interactions and take over the host cell machinery. Viral SLiMs may bind to host proteins (Davey, Trave et al. 2011, Mihalic, Simonetti et al. 2023), and folded viral proteins may bind to host SLiMs (Madhu, Davey et al. 2022, Mihalic, Benz et al. 2023). Both scenarios offer the possibility to inhibit viral infection by blocking the SLiM-binding pockets (Kruse, Benz et al. 2021, Mihalic, Benz et al. 2023, Simonetti, Nilsson et al. 2023). Finding and optimizing SLiM-based interactions between viral and human proteins thus offer potential strategies to develop antivirals.

Defining a SLiM requires both the identification of the binding peptide region and pinpointing key residues that confer affinity and specificity. Both can be accomplished by methods such as proteomic peptide phage display (ProP-PD) (Benz, Ali et al. 2022). However, in some cases, such analysis returns only limited sets of peptide ligands, preventing the identification of consensus motifs. Moreover, most approaches used assume that the binding determinants of the enriched peptides conform to one dominating motif consensus. Additional experiments, such as alanine scanning peptide arrays, point mutations, or structural analysis are subsequently required to define the binding determinant(s) for these peptides. Furthermore, motif variations and more subtle contributions of motif flanking residues are rarely captured by these approaches. During the last decade deep mutational scanning (DMS) has emerged as a powerful approach for gaining information on the functional effects of all possible mutations on binding (Fowler and Fields 2014). DMS can be used to evaluate how the binding between a protein and a given peptide is affected by replacing each amino acid in a peptide sequence with all other amino acids in a DMS library where the phenotype is linked to the genotype (e.g. by yeast display). Deep sequencing of the library before and after sorting/selection determines the relative abundance, and thereby the relative binding to the bait, of each sequence (Davey, Simonetti et al. 2023, Claussnitzer, Parikh et al. 2024). DMS has, for example, been utilized to explore motif-mediated interactions of PDZ and SH3 domains (Faure, Domingo et al. 2022), to characterize the LxxP docking motif for the yeast cyclin Cln2 (Bandyopadhyay, Bhaduri et al. 2020), to investigate the peptide binding of TRAF domains (Foight and Keating 2016) and to map antibody epitopes (Garrett, Itell et al. 2020).

In this study, we developed a phage display based DMS protocol for parallel analysis of distinct SLiM-based interactions. The method combines a designed oligonucleotide library, M13 peptide-phage display, and next-generation sequencing (NGS) (Fig. 1). We first evaluate the performance of phage based DMS by benchmarking the approach using a set of well-studied SLiM-based interactions (Benz, Ali et al. 2022, Kumar, Michael et al. 2024) and then applying it to a set of poorly characterized interactions involving peptides or folded domains from SARS-CoV-2 proteins (Kruse, Benz et al. 2021, Mihalic, Benz et al. 2023). We find that the DMS by phage display approach can be easily multiplexed and accurately determines binding specificities, pinpoints mutations that increase or decrease binding affinity, and provides directions for how to optimize the affinity of a given interaction. We demonstrate the utility of the strategy for engineering purposes by optimizing an antiviral peptide inhibitor binding to NSP9.

**Figure 1.**
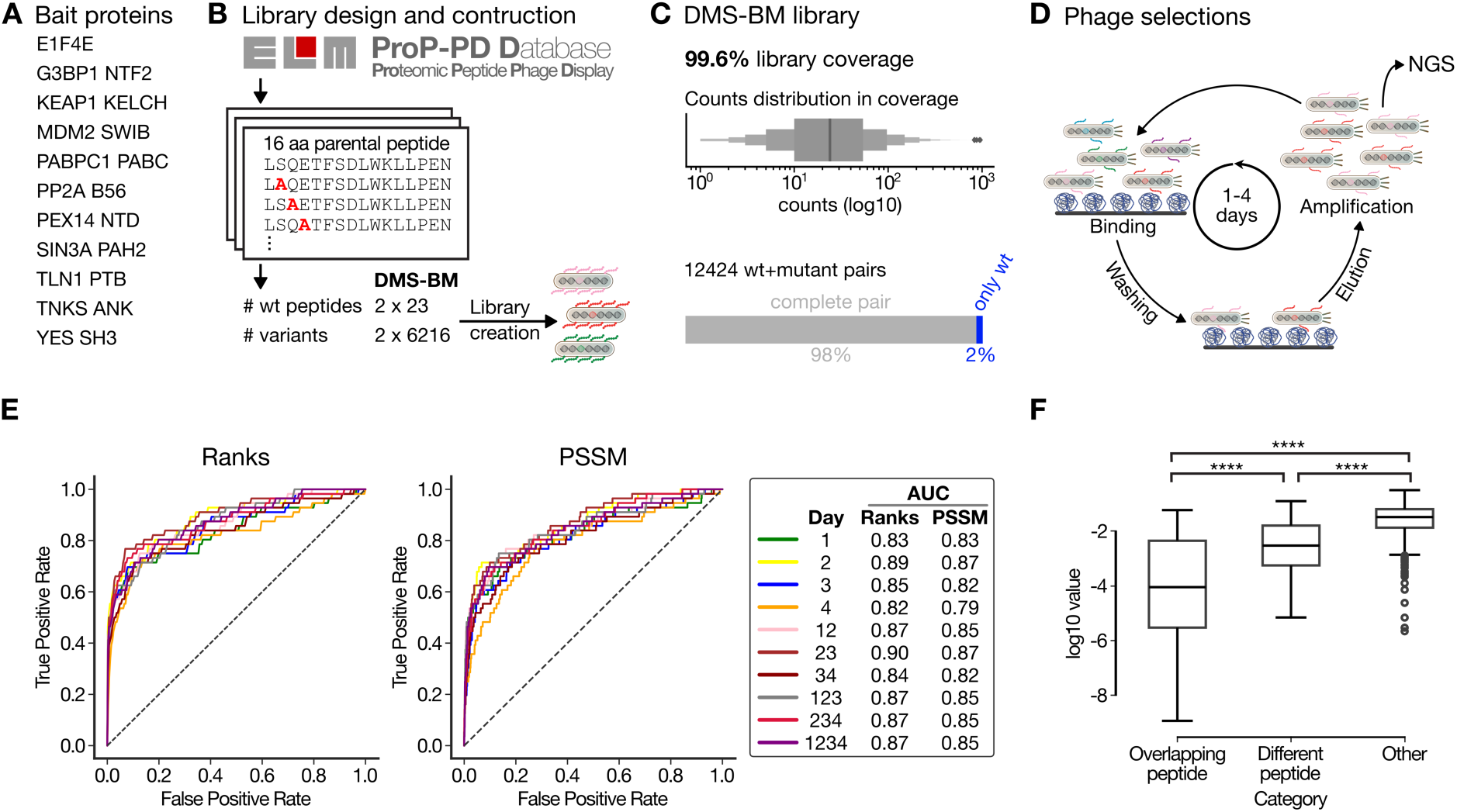
Overview of benchmarking DMS analysis. A) Bait protein domains selected and produced for benchmarking the DMS by phage protocol. B) Schematic of the design of the DMS-BM library. Overlapping parental peptides were selected based on data reported in the ELM database or in the ProP-PD portal, and used to generate the DMS phage library. C) Coverage of the DMS-BM phage library. D) Schematic of the selections against the DMS-BM library. The binding enriched phage pools were analyzed by NGS. E) ROC analysis of the correct identification of expected PSSMs from the DMS-BM selections as compared to the PSSMs generated based on peptide instances curated in the ELM database. F) Comparison of similarities of PSSMs generated based on results from overlapping peptides, from different peptides designed for the same bait, and PSSMs for unrelated baits. **** indicates p<0.0001.

## Results

### Design and construction of the DMS by phage benchmarking library

We designed a DMS benchmarking (DMS-BM) library (Table S1) to explore the effect of mutation on peptide binding to eleven different human bait protein domains (Fig. 1A), with distinct binding specificities (Table 1). Twenty-three peptide ligands were retrieved from the Eukaryotic Linear Motif (ELM) database (Kumar, Michael et al. 2024) or from the ProP-PD portal (Kliche, Garvanska et al. 2023) (Fig. 1B) The design included well studied interactions such as the p53 degron peptide binding to the SWIB domain of the E3 ubiquitin-protein ligase Mdm2 (MDM2) (Benz, Ali et al. 2022), and the cell division cycle-associated protein 2 (CDCA2) LxxIxE motif-containing peptide that binds to the B56 family protein phosphatase 2A (PP2A) regulatory subunit (Hertz, Kruse et al. 2016). Furthermore, we included peptides from USP10 and CAPRIN1 that both bind to the NTF2-like domain of G3BP1/2 but have distinct binding motifs (Song, Kuang et al. 2022, Schulte, Panas et al. 2023). Also, two distinct talin-1 (TLN1) PTB domain ligands were included that lack the canonical PTB binding NPxY motif (Benz, Ali et al. 2022). Each SLiM was tiled with two overlapping parental peptides shifted by two amino acids (14 amino acid overlap). Each position in the overlapping regions of the peptide pairs was subjected to in silico saturation mutagenesis (excluding cysteines for technical reasons) resulting in 12,432 peptides (Fig. 1B). The mutant peptide pool design was translated to oligonucleotides which were synthesized and genetically fused to the major coat protein P8 for multivalent display on the M13 phage. The sequence coverage of the constructed phage library was found to be 99.6%, with a balanced sequence representation (Fig. 1C).

**Table 1.** Overview of domains and peptides used for the design of the DMS-BM library, together with indication of the outcome of the analysis as evaluated by the sparsity score. Italic indicates residues which are found only in one of the two overlapping peptides included in the design. Bold residues indicate binding motif residues. The sparsity score ranges between 0-1 and a higher score indicates more informative DMS data.

| Domain | Peptide gene name | Peptide sequences | Sparsity of DMS results for peptide 1/peptide 2 |
| --- | --- | --- | --- |
| EIF4E<sub>1-217</sub> | EIF4EBP1 | <sup>50</sup>-*TRIIYDRKFL***MECRNSPV**<sub>-67</sub> | 0.53/0.44 |
|  | EIF4G1 | <sup>606</sup>-*LEEKKRYDREF***LLGFQFI**<sub>-623</sub> | 0.8/0.55 |
| G3BP1 NFT2<sub>1-139</sub> | CAPRIN1 | <sup>362</sup>-*LMAQMQGPYNFIQDSMLD*<sub>-379</sub> | 0.3/0.83 |
|  | USP10 | <sup>3</sup>-*LHSPQYIFGDF***SPDEFNQ**<sub>-20</sub> | 0.78/0.79 |
| KEAP1 KELCH<sub>321-609</sub> | NFE2L1 | <sup>226</sup>-*RNLLVDGET***GESFPAQVP**<sub>-243</sub> | 0.78/0.77 |
|  | SQSTM1 | <sup>342</sup>-*SSKEVD***PSTGELQSLQMP**<sub>-359</sub> | 0.66/0.68 |
| MDM2 SWIB<sub>17-125</sub> | KIAA1671 | <sup>600</sup>-*TPEDDRSFQTV***WATVFEH**<sub>-617</sub> | 0.67/0.81 |
|  | RNF115 | <sup>67</sup>-*TTTHFAELWGHLDHTMFF*<sub>-84</sub> | 0.79/0.83 |
|  | TP53 | <sup>14</sup>-*LSQETFSDL***WKLLPENNV**<sub>-31</sub> | 0.55/0.69 |
| PABPC1<br>PABC<sub>544-623</sub> | ATXN2 | 911-~~KSTLNPNAKEFNPRSF~~SQ<sub>-928</sub> | 0.6/0.74 |
|  | PAIP1 | 124-~~LMSKLSVNAPEFY~~PSGYS<sub>-141</sub> | 0.74/0.81 |
| PP2A B56<sub>1-486</sub> | AXIN1 | 235-~~SGYLPTLNEDEE~~WKCDQD<sub>-252</sub> | 0.71/0.77 |
|  | CDCA2 | 586-~~KKPLLSP~~IP~~ELPEVPE~~MT<sub>-603</sub> | 0.71/0.79 |
| PEX14 Pex14<sub>16-84</sub> | PEX5 | 108-~~GVADLALSENWAQ~~EFLAA<sub>-125</sub> | 0.67/0.82 |
|  |  | 238-~~AQAEQWAAEF~~IQQQGTSD<sub>-255</sub> | 0.7/0.85 |
| SIN3A PAH2<sub>295-383</sub> | KLF9 | 4-~~AAYMDFVAAQCL~~V~~SISNR~~<sub>-21</sub> | 0.57/0.28 |
|  | MXI1 | 4-~~VKMINVQRLLEAAEF~~L~~ER~~<sub>-21</sub><sup>27</sup> | 0.61/0.79 |
| TLN1 PTB<sub>309-401</sub> | PIP5K1C | 640-~~FPTDERSWVY~~SPLHYS<sub>-657</sub> | 0.84/0.92 |
|  | TPTE2 | 92-~~LADLIFTD~~SKLYIPLEYR<sub>-109</sub> | 0.71/0.65 |
| TNKS ANK<sub>174-649</sub> | AMOTL2 | 64-~~QVLQQATRQEPQG~~QEHQG<sub>-81</sub> | 0.82/0.8 |
|  | SH3BP2 | 408-~~PQLPHLQRSP~~PDGQSFRS<sub>-425</sub> | 0.82/0.84 |
| YES SH3<sub>90-152</sub> | BCAR1 | 627-~~DKTSSIQSRPL~~SPPKFT<sub>-644</sub> | 0.82/0.83 |
|  | CBL | 538-~~TLRDLPPPPPP~~DRPYSVG<sub>-555</sub> | 0.89/0.82 |

### DMS by phage correctly defines known SLiM consensuses

The DMS-BM library was used in phage selections against the eleven different bait proteins resulting in binding enriched phage pools (Table 1; Fig 1D; Table S2). The peptide-coding regions of enriched phage pools from day 1 to day 4 of selections were barcoded and analyzed by NGS. The resulting DNA sequences were translated into peptide sequences and position specific scoring matrices (PSSMs) were generated (Fig. S1) for each bait-peptide pair and each selection day. To evaluate the obtained results, we generated a benchmarking set of 234 PSSMs based on consensus aligned peptides from motif classes available in the ELM database (Kumar, Michael et al. 2024). The similarities between the ELM-based PSSMs and the PSSMs defined by the results of the DMS-BM selections were assessed for the results of each phage selection round (i.e. after 1, 2, 3 and 4 days of selection), as well as for combined selection round results (that is for round 1-2, 2-3, 3-4, 1-3 and 2-4). For each DMS-based PSSM, a similarity score *p*-value and a rank of the PSSM for its corresponding ELM-based PSSM in comparison to the remaining ELM classes screened were calculated. We performed a receiver operating characteristic (ROC) analysis of the PSSM similarity score *p*-value and the rank of the true positives ELM classes (True Positives) in relation to the remaining ELM classes (False Positives). The area under the curve (AUC) was calculated (Fig. 1E). The quality of the PSSMs varied by the selection day. The most informative results were obtained by combining the data of the second and third rounds of selections (AUC 0.9 for the rank, 0.87 for the PSSM similarity), and we thus used these data for the further analysis. Notably, the results of the selection day 2 were almost as informative by themselves based on the AUC (AUC 0.89 for the rank, 0.87 for the PSSM similarity), and the DMS analysis may thus be conducted using only two days of selection.

For each PSSM a sparsity score was calculated, and were found to be relatively high (Table 1). A high sparsity score (close to 1) indicates that sequencing data was obtained for a high proportion of the designed mutations. For the DMS We compared the PSSMs generated based on the selection results for a given bait-peptide pair against PSSMs for: (i) the same bait with the overlapping peptide; (ii) PSSMs for the same bait with a distinct peptide; and (iii) PSSMs for baits-peptide pairs from unrelated baits. As expected, we observed the highest PSSM similarity for overlapping peptides, followed by distinct peptides binding the same bait, and finally, limited similarity with other PSSMs in the dataset (Fig. 1F). The DMS-based PSSMs encode binding determinants that are similar to the binding determinants described in ELM. For example, the DMS of the p53 and RNF115 peptides binding to MDM2 resulted in the expected FxxxWxxL motif (Fig. 2A, B). Similarly, the DMS of the PP2A B56 binding peptides from AXIN1 and CDC2A resulted in a [LM]xx[ILV]xE motif, which closely resembles previously reported B56 binding LxxIxE motif (Fig. 2C,D) (Hertz, Kruse et al. 2016, Wu, Chen et al. 2017). For some baits, such as the G3BP1 NTF-like domain, we noted differences between the PSSMs generated using distinct model peptides. We used two distinct model peptides for G3BP1, one from USP10 and one from CAPRIN1, that are known binders of the same pocket but have distinct binding modes as shown by co-crystallization of the complexes (Song, Kuang et al. 2022, Schulte, Panas et al. 2023). The DMS analysis of the G3BP1 binding USP10 peptide (_3-_*LH*SPQYIFGDFSPDEF*NQ*_-20_) correctly identified its G3BP1 binding FG motif. The DMS analysis of CAPRIN1 peptide (_362-_*LM*AQMQGPYNFIQDSM*LD*_-379_) resulted in a distinct YxFI motif based on the averaged results of the two parental peptides. Notably, for the CAPRIN1_364-379_ peptide which generated the highest quality data (sparsity score 0.83) an extended YxFxxxSxL motif was obtained (Fig. S1). This is similar to the extended YNFIxxxxL G3BP binding motif previously observed for CAPRIN1 based on structural analysis (Schulte, Panas et al. 2023). As the terminal leucine is missing in the first CAPRIN1_362-377_ peptide the resulting averaged motif is truncated, suggesting that the frame of the peptides used may affect the motif observed. Nevertheless, the results demonstrate the potential of the DMS by phage display approach to reveal distinct motifs within different peptide-backbones.

**Figure 2.**
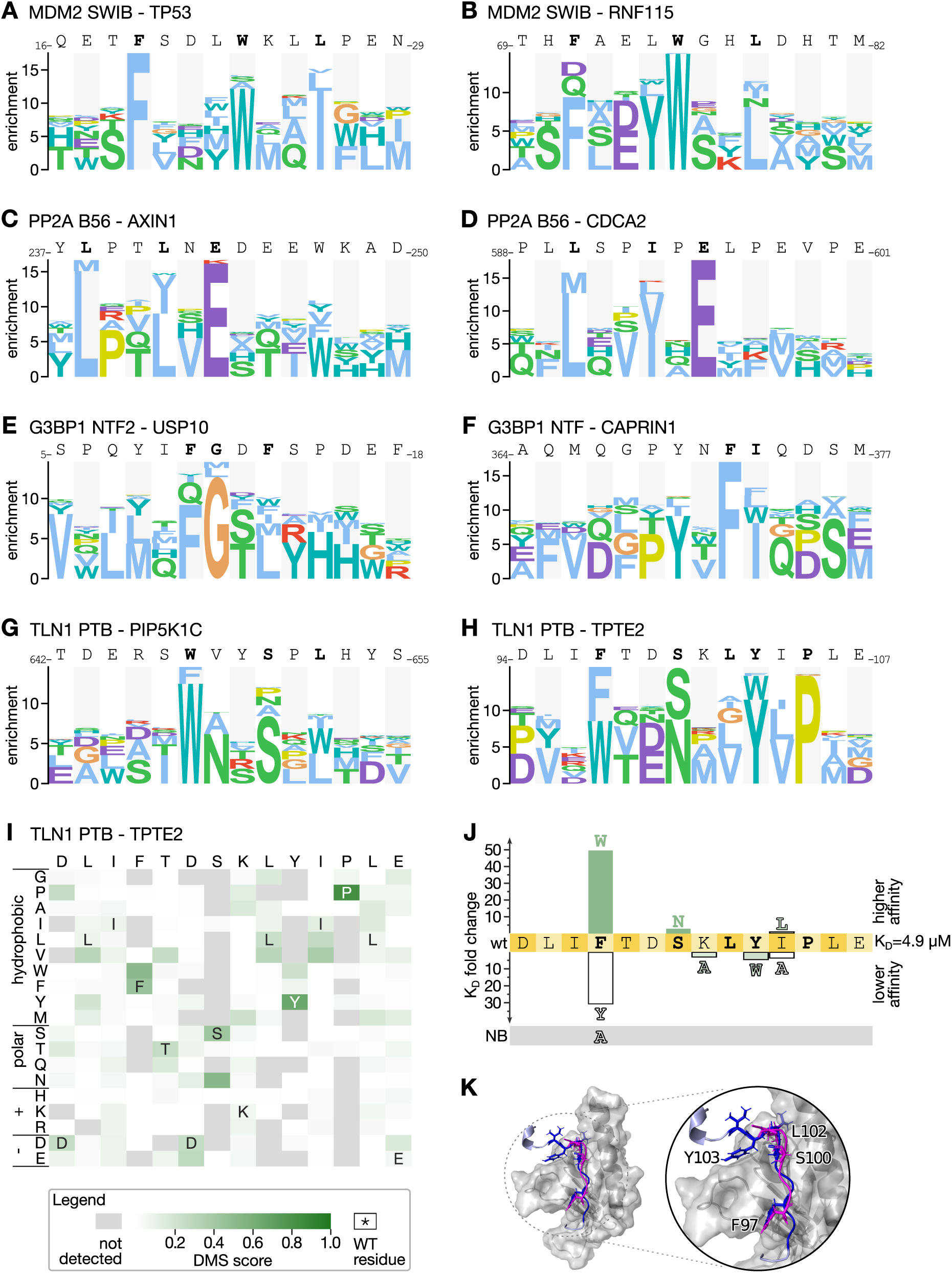
Examples of PSSMs generated by selections against the DMS-BM library together with validation of an extended TLN1 PTB binding motif in TPTE2. **A-H)** Representative examples of PSSMs generated for MDM2 (A, B), PP2A B56 (C, D), G3BP1 (E, F) and TLN1 (G, H). I) Heatmap representation of the PSSMs generated for the TPTE2 peptide binding to TLN1 PTB. J) Fold-change of affinities of TLN1 PTB binding TPTE2 peptides upon mutation as determined using fluorescence polarization-based affinity measurements. G) AlpaFold3 model of the TLN1 PTB-TPTE2 complex overlayed with the previously solved NMR structure of TLN1 PTB in complex with PIP5K1C (PDB id 2G35; peptide in magenta). The TPTE2 peptide is colored according to the pLDDT score

### DMS refines the talin-1 PTB binding determinants

For the atypical PTB domain of TLN1, we probed the two distinct peptides, a PIP5K1C peptide (_640-_ FPTDERSWVYSPLHYS_-657_) and a TPTE2 peptide (_92-_LADLIFTDSKLYIPLEYR_-109_). The DMS analysis revealed a common consensus motif in the two peptides, [WF]xxSxL, which in the TPTE2 peptide takes an extended form of [WF]xxSxLYxP (Fig. 2G, H). The TPTE2 peptide has a phenylalanine instead of a tryptophan at the first position of the motif, but the DMS results suggested that a tryptophan would be the preferred residue at the position. We therefore determined the affinities for the wild type TPTE2_92-107_ peptide, and its F97W, F97Y and F97A mutants using a fluorescence polarization (FP) based assay (Fig. S2, S3; Table S3). While the wild-type TPTE2_92-107_ peptide bound with a K_D_ value of 4.9 µM, the F97W mutant bound with 50-fold higher affinity (K_D_ = 0.1 µM; Fig. 2J). The F97Y mutation conferred instead a reduced affinity (K_D_=150 µM; 30-fold loss) while the F97A mutation resulted in loss of binding, highlighting the importance of the residue for binding. The DMS analysis further suggested that an asparagine is tolerated at the third position of the motif, and we found that a TPTE2_92-107_ S100N mutation conferred a minor increase in affinity (S100N; K_D_= 1.7 µM). We further explored the relevance of the putative extended motif in the TPTE2 peptide and found that mutation of Y103W (K_D_= 22 µM) and I104A (K_D_= 18 µM) reduced the affinity about 4-fold, while a conservative I104L mutation had minor effects, supporting that the TPTE2 exploits a longer motif, that is, that the motif-flanking region in the TPTE2 peptide contributes to binding. To gain further insight into how the TPTE2 peptide is bound by the TLN1 PTB domain we modelled the complex using AlphaFold3 (Fig. 2K) and overlayed it with the solved structure of the TLN1 PTB - PIP5K1C complex. The structural analysis showed that the [WF]xxSxL part of the two peptides bind in a similar way, with the [WF] at the first position docking into a hydrophobic pocket. While the PIP5K1C peptide loops out from the binding site, the C-terminal residues of TPTE2 peptide makes additional contacts with the domain. In particular, Y103 fits into a shallow pocket at the lid region. Taken together, based on the DMS analysis we define [WF]xx[SN]x[IL] as a general TLN1 PTB domain consensus motif, and show that the motif can be C-terminally extended.

### Exploring SLiM-based host-virus interactions by DMS by phage display

Having benchmarked the DMS by phage display protocol and showed its potential for uncovering novel details of well-studied interactions we next applied the approach to less explored host-pathogen interactions. We designed a second DMS library (Table S1), termed DMS-CoV, based on eight viral peptides binding to five human bait protein domains and eleven human peptides binding to five SARS-CoV-2 bait protein domains (Table 2). The studied interactions included among others: two viral peptides binding to the G3BP1/2 NTF2-like domain, and human ligands of the globular domains of NSP3 and of NSP9 (Fig. 3A). The interactions were previously found through proteomic peptide phage display (Kruse, Benz et al. 2021, Mihalic, Benz et al. 2023) or predicted based on consensus binding motif (i.e. the SH3 binding PxxP motif in the SARS-CoV-2 N binding to ABL1 SH3 domain). NGS analysis confirmed that 96.5% of the designed oligonucleotides were represented in the constructed library (Fig. 3B). While the sequence coverage was high, there were systematic deviations such that the SH3 binding _359-_DAYKTFPPTEPKKDKKKK_-376_ peptide from the N protein and its variants were depleted in the constructed phage library, possibly due to the lysine-rich peptide interfering with phage virion assembly. Thus, the coverage of the DMS-CoV library at the peptide level was lower than of the DMS-BM library (Fig. 3C). Nevertheless, we used the DMS CoV library in selections against the defined bait collection. The results obtained using the DMS-CoV library were less informative than the results obtained using DMS-BM library (Fig. 3D), partially due to the lower coverage but likely also due to the fact that the interactions probed were of lower affinity (Mihalic, Benz et al. 2023) and due to some traits of the motifs as described below. In several cases, only one of the two overlapping parental peptides returned sufficient data, which may indicate that parts of the motifs were truncated in the shifted peptides. For example, for the EZR FERM domain the analysis correctly identified the YxΦ motif in the N-terminal part of the envelope (E) protein peptide (Fig. 3E). The motif is lost in the shifted peptide, which explains the lack of information obtained for the second parental peptide tiling the region. Finally, the viral USP7 MATH domain ligands included in the design failed to be enriched in the selections as they were outcompeted by peptides from the MBOAT1 and AZIN2, which turned out to contain uncharacterized USP7 binding motifs (Fig. S4). The results highlight that factors such as the affinity of the interactions probed and the position of the motif in the peptide should be considered when designing libraries for multiplexed DMS by phage display experiments.

**Figure 3.**
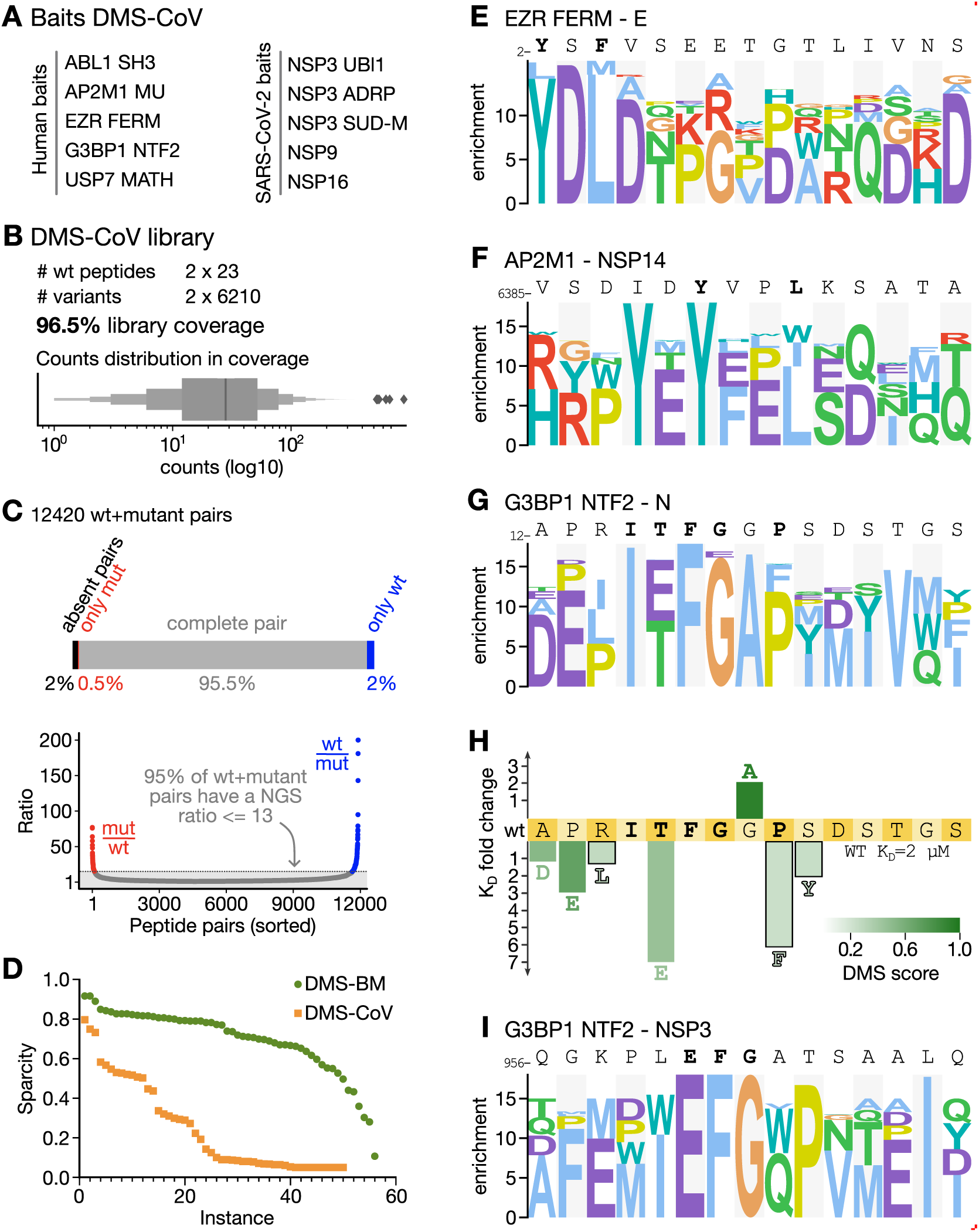
DMS-CoV library quality and results for selections against human bait proteins. A) Bait proteins selected for the DMS-CoV analysis. B) DMS-CoV phage library design parameters, coverage and count distribution on peptide level .C) DMS-CoV library coverage on the peptide/wild-type pair level. D) Sparsity of DMS-CoV selection results as compared to the DMS-BM results. A low sparsity score (y-axis) indicates that sequencing data is missing for many mutations and amino acid positions for a given parent peptide (instance, x-axis). E-G, I) Representative PSSMs generated for viral peptides binding to the human bait protein domains EZR FERM (E) AP2 M1 (F), and G3BP1 NTF2 (G, I). H) Fold-change of affinities of G3BP1 NTF2-binding N peptides upon mutation.

**Table 2.**
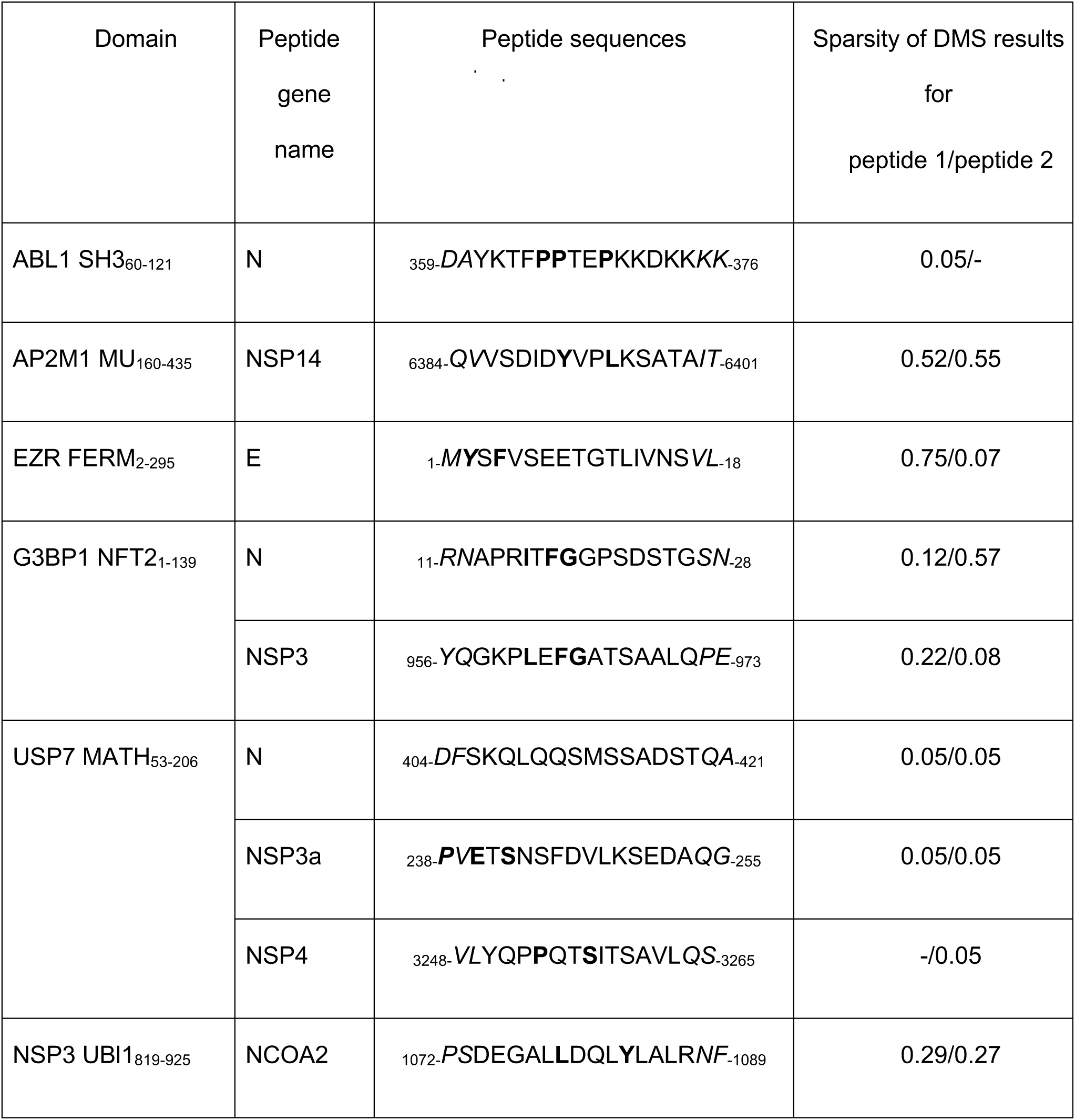
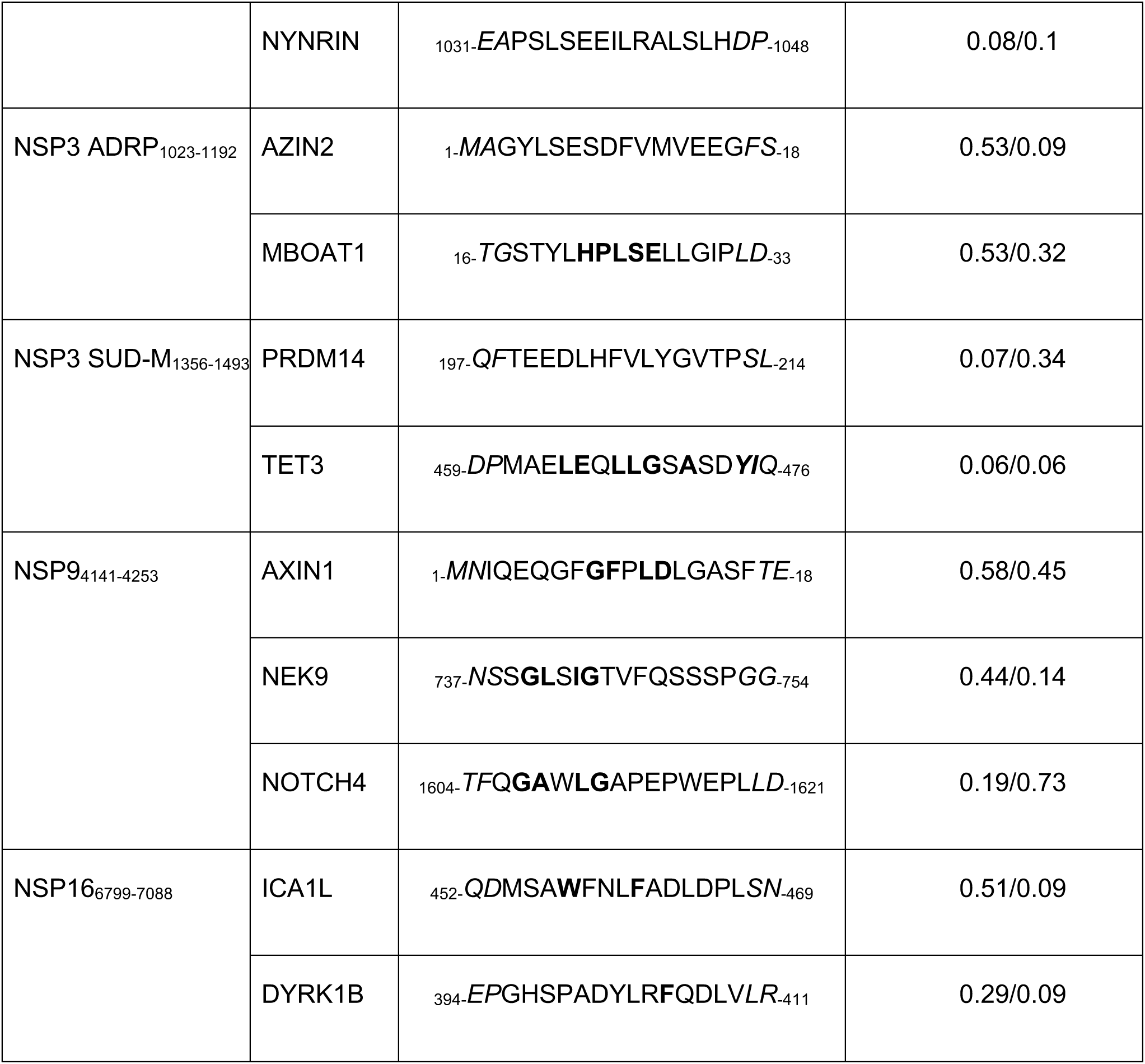
Overview of domains and peptides used for the design of the DMS-CoV library, together with indication of the outcome of the analysis as evaluated by the sparsity score. Italic indicates residues which are found only in one of the two overlapping peptides included in the design. Bold residues indicate binding motif residue based on consensus motifs or previous alanine scanning SPOT array analysis. The sparsity score ranges between 0-1 and a higher score indicates more informative DMS data.

### DMS results for SARS-CoV-2 peptides binding to the NTF-like domains of G3BP1

Among the viral peptides binding to human proteins, we found two interactions particularly interesting. Firstly, our analysis confirmed the expected YxxL AP2M1 binding motif in the probed peptide from NSP14 (Fig. 3D), but also suggested that an additional AP2M1 motif can emerge in the peptide upon an isoleucine to tyrosine substitution (YxxV), resulting in two potentially overlapping AP1M1 binding sites in the same peptide. Secondly, the DMS analysis correctly showed that the two viral G3BP1 binding peptides, N_11-28_ (K_D_ = 2.3 µM) and NSP3_956-973_ (61 µM) (Kruse, Benz et al. 2021) share an (E/T)FG motif (Fig. 3G, I), similar to the FG motif found in USP10 (Fig. 2). A T16E mutation in the N peptide conferred a 4-fold loss of affinity, partly explaining the higher affinity of the N peptide for G3BP1 as compared to the NSP3 peptide (N_12-26_ T16E = 16 µM; Fig. 3H). In addition, the DMS results for the N peptide suggested that the interaction is supported by motif flanking residues (IT<u>FG</u>xP), which is consistent with the binding determinants (ITFG) resolved through co-crystallization of the G3BP1 NTF-N peptide complex (Biswal, Lu et al. 2022). The results further indicated that the proline contributes to binding, as a P20F mutation conferred a 6-fold loss of affinity for N_12-26_. The DMS results further suggested that the affinity of the N peptide for G3BP1 could be improved by mutating a glycine in a wild-card position to alanine, and affinity measurements confirmed that the G19A mutation conferred a two-fold increase in affinity (K_D_ wildtype N_12-26_ = 2.3 µM; N_12-26_ G19A = 1.1 µM). Other mutations tested in the flanking residues conferred moderate or minor losses of affinity (Fig. 3H).

### Refining the motifs in human peptides binding to protein domains from SARS-CoV2 proteins

We next turn to the analysis of human peptides binding to viral protein domains expressed by the SARS-CoV-2 genome. We previously uncovered peptide-based interactions of three NSP3 domains, NSP9, and NSP16 (Mihalic, Benz et al. 2023), which were further explored here.

#### NSP3

The large multidomain protein NSP3 has several peptide binding domains including NSP3 ADRP, NSP3 UBl1, and NSP3 SUD-M. For NSP3 ADRP we tested two peptides previously identified as binders (AZIN2_1-18_ and MBOAT1_16-33_). A previous alanine scanning SPOT array analysis suggested the core motif in the MBOAT peptide to be HPLSE, and the current DMS analysis confirmed that this is a critical region for binding (Fig. 4A, D). Based on the DMS results we attempted to improve the affinity of the MBOAT1 peptide for NSP3 ADRP by a set of point mutations in the flanking regions, but the mutations resulted in minor (2-3 fold) losses of affinity in comparison to the wild-type MBOAT1_16-31_ peptide (K_D_ = 49 μM) (Fig 3D; Fig. S3; Table S3).

**Figure 4.**
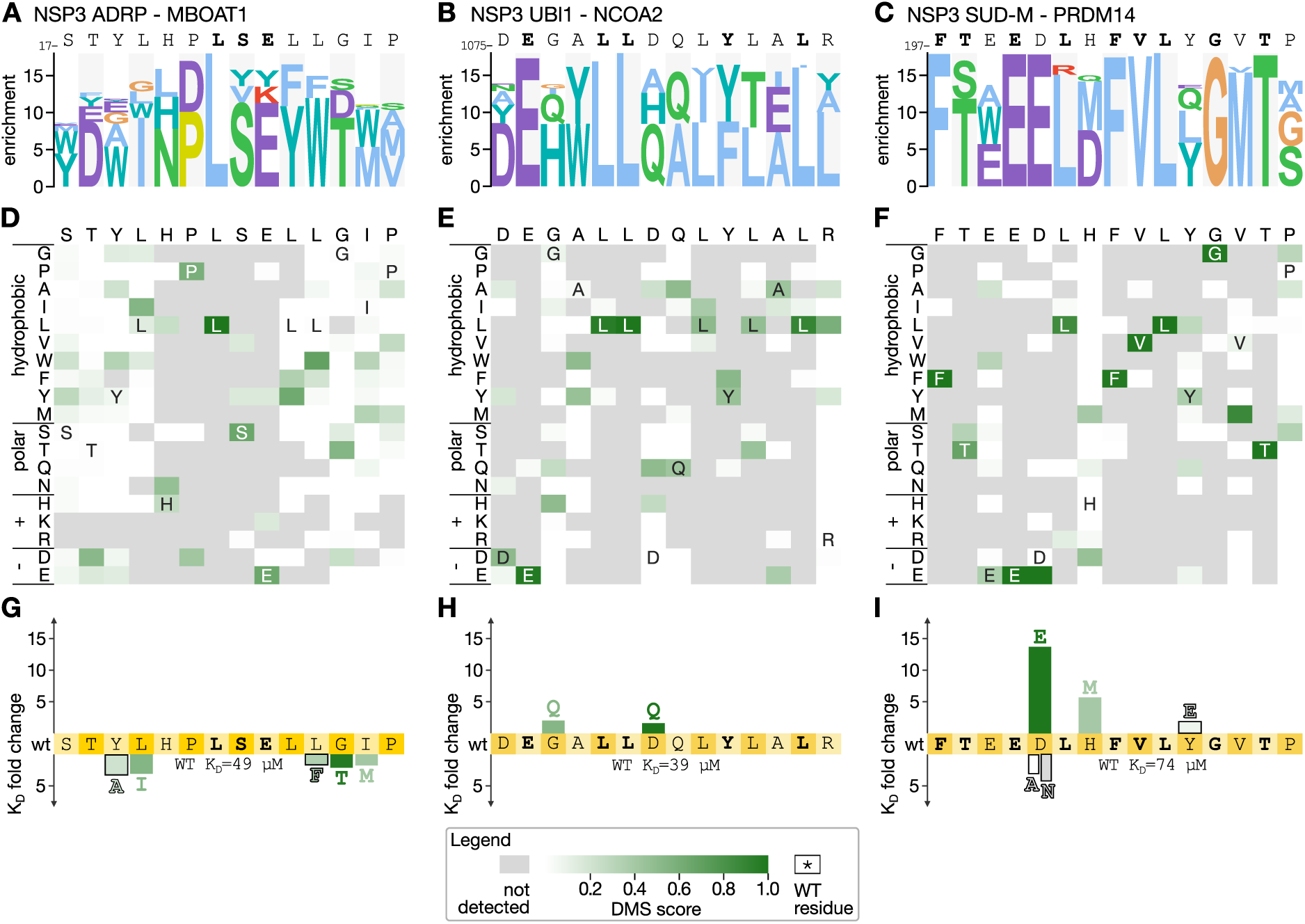
DMS analysis of peptides binding to the ADRP, UBl1 and SUD-M domain of NSP3. PSSM (A-C) and heatmap (D-F) representations of the DMS data for the MBOAT1 peptide binding to NSP3 ADRP (A,D), the NCOA2 peptide binding to NSP3 UBl1 (B, E) and the PDRM14 peptide binding to NSP3 SUD-M (C, F). G-I) Fold-change of affinities upon mutation of the respective wild-type peptide binding to NSP3 ADRP (G), UBl1 (H) and SUD-M (I).

For the NSP3 UBl1 domain, we also subjected two peptides for DMS analysis (NYNRIN_1031-1048_ and NCOA2_1072-1089_), of which NCOA2 is the higher affinity ligand (Mihalic, Benz et al. 2023). Consistent with its higher affinity, the most informative results were obtained for the NCOA2_1072-1089_ peptide (Fig. 4B, E) that converged on an extended ExxLLxxxYxxL motif. The extended motif partially matches the LxxxY motif previously suggested based on SPOT array alanine scanning (Mihalic, Benz et al. 2023). In an attempt to increase the affinity of the interaction we tested two mutations (G1077Q and D1081Q) and evaluated their effects on binding. Each of the mutations conferred minor increases in affinity in comparison to the wild-type peptide (Fig. 4H; K_D_ = 19 and 24 μM for G1077Q and D1081Q, respectively, in comparison to 39 μM for wild-type NCOA2_1073-1088_).

For NSP3 SUD-M we tested the two model peptides, PRDM14_197-214_ and TET3_459-476_, of which the PRMD14_197-214_ peptide (_197-_*QF*TEEDLHFVLYGVTP*SL*_-214_) is the higher affinity ligand (Mihalic, Benz et al. 2023). Consistently, the DMS selection was dominated by the PRDM14 peptide and its variants (Fig. 4G, F). The DMS analysis suggested that the peptide contains an extended <u>F</u>xx<u>E</u>xLx<u>F</u>V<u>L</u>x<u>G</u>x<u>T</u> motif, which is similar to the previous results obtained through SPOT array alanine scanning (underlined; (Mihalic, Benz et al. 2023). We designed two mutations to improve the affinity of the interaction (D201E, H203M), and also tested a Y207E thought to be largely neutral to binding, and, as a control, included mutations that were expected to decrease the affinity (D201A, D201N). Affinity measurements revealed that the conservative D201E mutation had the most beneficial impact on binding (K_D_ = 5.4 μM, 17-fold increase in affinity compared to wild-type peptide Kd of 74 μM), followed by the H203M (K_D_ = 13 μM). The Y207E mutation also led to a minor improvement of affinity (K_D_ = 40 μM) (Fig. 4I; Fig. S3; Table S3). As expected, the D201A and D201N mutations conferred reduced affinity.

In summary, the DMS analysis of the peptides binding to the NSP3 domains validated their key residues and pinpointed ways to improve their affinities, in particular for the NSP3 SUD-M domain.

#### NSP9

NSP9 is a component of the SARS-CoV-2 5’ mRNA capping machinery. We previously reported that it binds to a large number of peptides from human proteins containing a GΦxΦ[GD] motif, where Φ is a hydrophobic amino acid (Mihalic, Benz et al. 2023). Here, we probed its binding to three peptides, from AXIN1, NEK9 and NOTCH4, respectively, where analysis of AXIN1 and NOTCH4 peptides produced the most robust data. The DMS analysis of the AXIN1 peptide resulted in a G[LVF]x[IL]D motif (Fig. 5A), while the NOTCH4 peptide instead revealed a similar yet distinct GxWLG motif (Fig. 5B,C). Affinity measurements of the wild-type and mutant NOTCH4 peptides and NSP9 showed that a glycine to aspartic acid (G1611D; K_D_ = 160 μM), or a or a glycine to proline (G1611P; K_D_ = 410 μM) substitution at the last position of the motif conferred losses of affinity (2 to 5-fold) as compared to the affinity for the wild-type NOTCH4 peptide (K_D_ = 72 μM), thus supporting the motif variations between the two model peptides (Fig. S3; Table S3).

**Figure 5.**
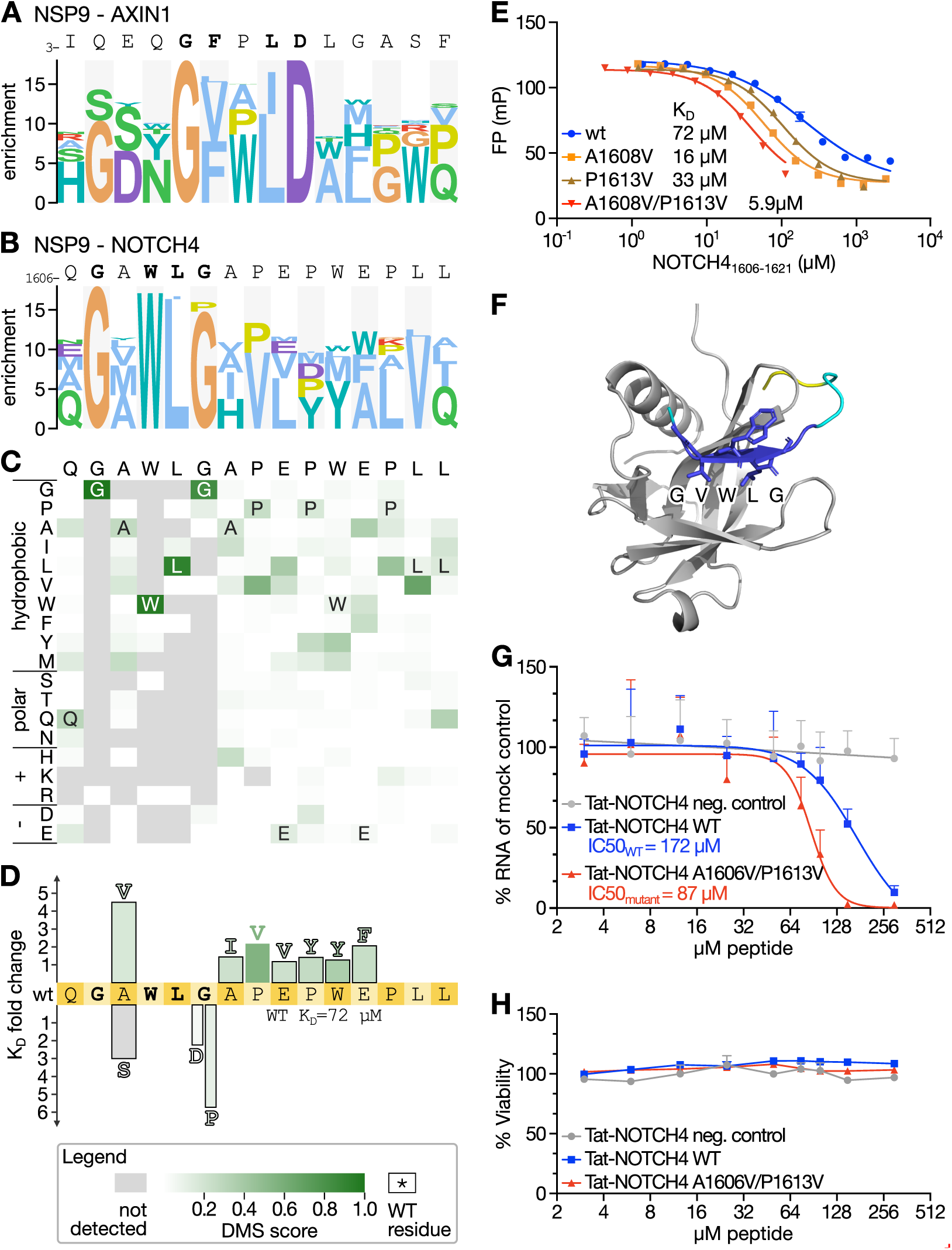
DMS analysis of NSP9 binding peptides guides the engineering of more potent antiviral inhibitors. A) PSSM representation of the DMS results of the Nsp9 binding AXIN1 peptide. B, C) PSSM and heat map representation of the of the DMS results of the NSP9 binding NOTCH4 peptide. D) Fold-change of affinities upon point mutation of the NOTCH4 peptide. E) FP-based affinity determinations of wt, single (A1603V and P1613V) and double mutants of the NOTCH4 peptide binding to NSP9 (A1603V/P1613V). F) AlphaFold3 model of the complex of NSP9 and the NOTCH4 A1603/P1613V peptide. Peptide coloring is according to the pLLDT score (deep blue = high confidence). G) Evaluation of the antiviral effect of cell-permeable Tat-tagged variants of the NOTCH4 peptides in VeroE6 cells. VeroE6 cells were infected with SARS-CoV-2 (multiplicity of infection: 0.5) and 8 hours post-infection viral RNA was quantified using qPCR. Viral RNA was normalized to the RNA levels in mock-treated cells and presented as % RNA of control. Data are cumulative of two independent experiments done in triplicates (N = 6). H) Unaffected cell viability upon treatment with the Tat-tagged peptides. Cellular viability after peptide treatment was measured using Celltitre Glo. Data are cumulative of two independent experiments done in triplicates (N = 6).

As the NSP9 binding NOTCH4 peptide has previously been shown to have antiviral effect (Mihalic, Benz et al. 2023) we further attempted to generate a higher affinity NOTCH4 ligand based on the DMS data. We tested seven mutations and found each of them to confer minor increases in affinities, with a conservative A1608V mutation at the second position of the motif having the largest effect (A1608V, K_D_= 16 µM versus K_D_= 72 µM for the wild-type; Fig. 5D, 5E). As a negative control, we tested a S1608V mutation, which as expected led to a decreased affinity (K_D_= 220 µM). We combined the A1608V mutation with a P1613V mutation (K_D_= 33 µM) into a double mutant, which resulted in a further increase in affinity (A1608V/P1613V K_D_= 5.9 µM). Having generated a higher affinity NSP9 ligand we used AlphaFold3 (Abramson, Adler et al. 2024) to model the complex. In contrast to the wild-type peptide, the NOTCH4 A1608V/P1613V variant was confidently docked, with the model suggesting that the GVWLG part of the peptide binds through beta-strand addition (Fig. 5F). The proposed NSP9 binding region coincides with residues previously mapped by NMR to be perturbed by peptide binding (Mihalic, Benz et al. 2023). Thus, the previously NMR-mapped binding site and the AlphaFold3-based model confidently pinpoint the peptide binding region on NSP9.

Finally, we tested the antiviral activity of the affinity matured NOTCH4 A1608V/P1613V peptide by fusing it to a cell-penetrating Tat-tag and evaluated its antiviral effect in comparison to the Tat-tagged NOTCH4 wild-type peptide. VeroE6 cells were treated with the Tat-tagged peptides and infected with SARS-CoV-2 (multiplicity of infection: 0.5). The viral RNA was quantified 8 hours post infection using qPCR. The analysis showed that the Tat-tagged NOTCH4 A1608V/P1613V peptide is a more potent antiviral inhibitor consistent than the wild-type peptide, consistent with the higher affinity (Fig. 5G), while not having any effect on cell-viability (Fig. 5H).

## Discussion

In this study we outline and benchmark a multiplexed DMS by peptide-phage display protocol. Through benchmarking the results against a set of well-defined motifs reported in ELM (Kumar, Michael et al. 2024) we show that DMS by phage display is an efficient approach for defining interaction motifs. The phage display-derived DMS data can identify the key residues in the motif and define preferred amino acids in these position in the context of the chosen peptide. In addition to consensus discovery, we find that the DMS analysis provides useful information on the contribution of motif flanking residues, and pinpoints variations of the motifs that are not captured by the general motif descriptions or from consensus motifs generated by aligning cohorts of binding peptides. For example, we highlight the case of the TLN1 PTB domain, for which we defined a general consensus motif ([WF]xx[SN]x[IL]), which can be supported by a C-terminal extension ([WF]xx[SN]x[IL]<u>Y</u>x<u>P</u>). The two similar yet distinct motifs found in the G3BP1-binding peptides from USP10 and CAPRIN1 support this point.

We further applied the “DMS by phage display” protocol on less explored SLiM-based host-virus interactions. This analysis turned out to be more challenging, likely due to the lower affinities of the interactions. Nevertheless, the DMS analysis confirmed and substantiated the previously described motifs binding to the SARS-CoV-2 domains NSP3 UBl1, NSP3 ADRP, NSP9 and NSP16. The data also revealed detailed motif variations in the peptides binding to NSP9. The results support that the general NSP9 binding motif is GΦxΦ[GD], and that the NOTCH4 peptide has a similar yet distinct motif (GxWLG). The two variant motifs dock to the same site based on AlphaFold3 modelling (Fig. S5), and the motif variation appears to be caused by the requirements posed by the need to accommodate a bulky tryptophane in the NOTCH4 upon beta-strand addition. Moreover, we showcase how DMS by phage display can be used to increase the affinity of peptide ligands, and that the affinity-matured NSP9 binding peptide has increased antiviral activity. Consequently, DMS by phage display can be used both to identify the determinants of host-virus protein-protein interactions and to increase the affinity of peptide ligands as a part of peptide-based inhibitor development.

In summary, we find that a strength of DMS by phage display is the scalability as it can easily be performed in parallel for multiple peptides binding to various protein domains. Given the multiplexing possibilities, we envision the integration of DMS into a workflow where a limited set of ligands has been identified for several different SLiM-binding protein domains. A limitation of the approach is that it does not perform well for low affinity interactions (e.g. with K_D_ values above 100 µM), that care needs to be taken when designing the library (e.g. choice of model peptides). When combining multiple DMS analyses into one experiment there is also the risk of unexpected competition between different ligands targeting the same pocket. With these limitations in mind, we conclude that DMS by peptide-phage display can be applied to obtain information on binding determinants for multiple proteins in parallel and pinpoint the divergent affinity determinants in distinct peptide backgrounds. DMS by peptide-phage display thus represent a viable addition to the toolbox for exploration of SLiM-based interactions.

## Material and Methods

### Library design

The DMS-BM and DMS-CoV phage libraries were designed based on previously reported ligands (Kruse, Benz et al. 2021, Benz, Ali et al. 2022, Mihalic, Benz et al. 2023, Kumar, Michael et al. 2024).. Each wild-type peptide was tiled by two overlapping peptides shifted by 2 amino acids. All wt peptides were mutated on all overlapping positions to all-natural amino acids except cysteine. The peptides were reverse translated to oligonucleotides optimized for *E. coli* expression and flanking regions for library creation were added (5’ CAGCCTCTTCATCTGGC and 3’ GGTGGAGGATCCGGAG).

### Phage library constructions

The oligonucleotides (GenScript) were PCR amplified with Phusion PCR Master Mix (Fisher Scientific) using 90 sec 98°C initial denaturation; 18 cycles of 15 sec 98°C denaturation, 15 sec 55-58°C annealing, and 15 sec 72°C elongation; and 5 min 72°C final elongation. The PCR products were purified using the MinElute PCR Purification Kit (Qiagen). The PCR-amplified oligonucleotides (0.6 μg) were 5’ phosphorylated with 20 units of T4 polynucleotide kinase (Fisher Scientific) at 37°C for 1 h in 1x TM buffer (10 mM MgCl_2_, 50 mM Tris-HCl, pH 7.5) supplemented with 5 mM dithiothreitol (DTT) and 1 mM adenosine triphosphate (ATP). Following 5 minutes of cooling on ice, the oligonucleotides were annealed to 10 μg of dU-ssDNA phagemid (90°C for 3 min, 50°C for 3 min, and 20°C for 5 min) in TM buffer. DNA polymerization and ligation were initiated by adding 10 μL 10 mM ATP, 10 μL 10 mM dNTP, 15 μL 100 mM DTT, 30 Weiss units of T4 DNA ligase (Fisher Scientific), and 30 units of T7 DNA polymerase (Fisher Scientific), followed by incubation at 20°C for 16 h. The reaction was stopped by three freeze-thawing cycles. Remaining wild-type dU-ssDNA was digested by incubating with 5 μL FastDigest SmaI (Fisher Scientific, 37°C, 30 min). dsDNA was purified using the QIAquick PCR & Gel Cleanup Kit (Qiagen). The dsDNA phagemid library was electroporated into *E. coli* SS320 cells (Lucigen) pre-infected with M13KO7 helper phages (ThermoFisher). Electroporated cells were rescued in 25 mL pre-warmed super optimal broth (SOC) medium (0.5 w/v% yeast extract, 2 w/v% tryptone, 10 mM NaCl, 2.5 mM KCl, 10 mM MgCl_2_, 10 mM MgSO_4_, and 20 mM glucose, pH = 7.0) and incubated at 37°C for 30 min. The phage-producing bacteria were grown overnight (±18 h) in 0.5 L 2YT medium (1 w/v% yeast extract, 1.6 w/v% tryptone, and 0.5 w/v% NaCl) at 37°C, and then harvested. Phage libraries were stored at -80°C in 10 v/v% glycerol.

### Bait expression and purification

The pETM33 (EMBL), PH1003 (Sidhu Lab), pET42a(+) (EMD Biosciences), or pGEX-4T1 (GenScript) vectors containing cDNA encoding bait proteins (Table S2) were used to express GST-tagged baits. Overnight cultures (2YT, 37°C, 200 rpm, 18 h) of *E. coli* Bl21 DE3 gold cells (Agilent) transformed with the appropriate vector were used to inoculate 500 mL 2YT (supplemented with Kan (50 μg/mL) or Carb (100 μg/mL)), followed by incubation (37°C, 200 rpm). Protein production was induced at OD_600_ 0.6-0.8, with 1 mM IPTG for 18-20 h at 18 °C, 200 rpm. The bacteria were pelleted (5000 xg, 5-7 min) and stored at -20°C. Bacteria were dissolved in lysis buffer (phosphate buffered saline (PBS, 37 mM NaCl, 2.7 mM KCl, 8 mM Na_2_HPO_4_, 2 mM KH_2_PO_4_), pH 7.4, 1% Triton X-100, 10 μg/mL DNase I, 5 mM MgCl_2_, lysozyme (Thermo Scientific), cOmplete Mini, EDTA-free, Protease Inhibitor tablet (Roche, 1 tablet/10mL)), incubated at 4°C, for 1 h and sonicated (2 sec pulse, 2 sec pause for 20 sec). Cell debris were removed (16.000 xg, 4°C, 1 h). The supernatant was incubated with Glutathione (GSH) Sepharose 4 Fast Flow beads (Cytiva) (4°C, agitation, 1 h). Protein purities were confirmed through SDS-PAGE (BioRad Mini-PROTEAN TGX Stain-Free Precast gels, 200 V, 30 min). Purified bait proteins were flash frozen using liquid nitrogen in 16 v/v% glycerol and stored at -80°C until further use.

### Phage selections

10 μg of GST-tagged bait proteins or GST (negative control) in 100 μL PBS were immobilized in Nunc MaxiSorp flat-bottom 96-well plates (ThermoFisher Scientific, Cat: 44-2404-21) for 18 h at 4°C. Wells were blocked with 200 μL 0.5% bovine serum albumin (BSA) in PBS for 1 at 4°C under gentle agitation. GST-coated wells were washed four times with 200 μL PT (PBS + 0.05 v/v% Tween 20) and (naïve) phage library (10^11^ phages, 100 μL in PBS) was added to each GST-coated well. Following incubation (4°C, gentle agitation, 1 h), the phage library was transferred to blocked and washed bait-protein-coated wells. After 2 h of incubation at 4°C, unbound phages were removed by five 200 μL PT washes. Bound phages were eluted with 100 μL log-phase *E. coli* OmniMAX cells (cultured in 2YT medium supplemented with 10 mg/mL tetracycline) for 30 min at 37°C under gentle agitation. 10^9^ M13KO7 helper phages (ThermoFisher) were added to each well and allowed to infect bacteria for 45 min at 37°C. The hyper infected bacteria were transferred to 1 mL 2YT supplemented with 30 μg/mL Kan, 100 μg/mL Carb, and 0.3 mM IPTG and incubated over-night (37°C, 200 rpm, ±18 h). Bacteria were pelleted (2000 xg, 4°C, 10 min) and the phage supernatants were transferred to a fresh 96-deep-well plate, pH adjusted by adding 1/10 volume 10x PBS and heat-inactivated through incubation at 65°C for 10 min. The phage pools were used for the next day of selection.

### Phage Pool ELISA

Proteins (10 μg) in PBS (100 μL/well) were coated in a Nunc MaxiSorp flat-bottom 96-well plates for18 h at 4°C under gentle agitation. Wells were blocked with 200 μL 0.5% BSA in PBS (4°C, 1 h). Phages (100 μL) were allowed to bind to the bait protein- or GST-coated wells for 1h at 4°C. Unbound phages were washed away with 4x 200 μL PT and 100 μL HRP-conjugated anti-M13 bacteriophage antibody was added (Sino Biological Inc, Cat: 11973-MM05T-H, 1:5000 diluted in 0.5% BSA in PT). Following a 1 h incubation at 4°C, wells were washed four times with 200 μL PT and once with 200 μL PBS. 100 μL TMB substrate (Seracare, Cat: 5120-0047) was used to detect the bound antibody and the enzymatic reaction was stopped through addition of 100 μL 0.6 M sulfuric acid (H_2_SO_4_). The absorbance at 450 nm was measured with a SpectraMax iD5 microplate reader (Molecular Devices).

### Sample preparation for NGS and data analysis

Peptide-coding regions of were amplified and barcoded using Phusion PCR Master Mix (Fisher Scientific) for 22 cycles. PCR products (25 μL) were normalized using Mag-Bind Total Pure NGS magnetic beads (Omega Bio-Tek, Cat: M1378). Normalized PCR products (10 μL) were pooled and purified using gel purification. DNA was eluted with 30 μL TE buffer (10 mM Tris-HCl, 1 mM EDTA, pH 7.5). The amplicon pool was sent for NGS (Illumina MiSeq v3, 1x150bp read setup, 20% PhiX, performed by the NGS-NGI SciLifeLab facility). The raw NGS data was demultiplexed and translated to peptide sequences using custom Python scripts.

### ELM instance specificity determinant dataset

A dataset of PSSMs encoding motif class specificity determinants was created from the motif instances in the ELM database (Kumar, Michael et al. 2024). For each SLiM class, peptides were extracted and aligned using the ELM-defined class consensus, alignments were converted to a PSSM using the PSSMSearch web application (Krystkowiak, Manguy et al. 2018) with default parameters and the frequency PSSM scoring method, resulting in 234 PSSMs. Each bait-peptide pair screened in the DMS-BM analysis was annotated with a corresponding ELM class.

### Specificity determinant comparison

The similarities between the specificity determinants resulting from the DMS-BM screens and the expected specificity determinants were quantified using PSSM-PSSM comparison. The comparison is performed by sliding two PSSMs across each other and calculating the similarity of each comparison window. The similarity of each corresponding column in the window was calculated using Pearson’s correlation. The importance of the amino acid position was also calculated for each column using the Gini Coefficient, a measure of statistical dispersion that calculates the inequality among values. The importance-weighted similarity score (ISW) is then calculated using the following equation:

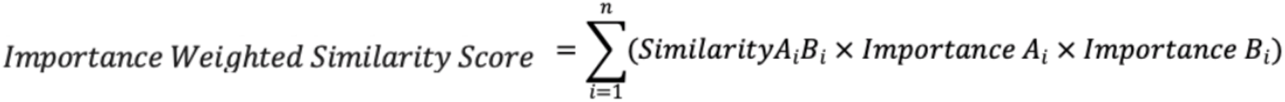

***Equation 2:*** *ISW calculation,* where n is the number of positions in the motif alignment, A_i_ is position *i* in PSSM_A_ and B_i_ is position *i* in the PSSM_B_.

The probability of the observed importance-weighted similarity score between two columns was calculated using a randomization approach based on the comparison of random PSSM columns. A sample of 100,000 randomly selected column pairs between the two PSSM datasets were compared, and the distribution of importance weighted similarity score was calculated. The likelihood of seeing the observed importance-weighted similarity score by chance, *p^ISW^*, was defined based on the distribution of the importance-weighted similarity score of the randomly paired PSSM columns. The probability of the window, *p^window^*, was calculated as the product of the pairwise column *p^ISW^* probabilities from the window. The *p^window^* score was normalized to correct for the number of comparisons performed for the window using uniform product distribution correction to define the *p^window_corrected^* probability. After all the comparison windows were scored, the highest-scoring pair of windows was returned as the aligned specificity determinants and the *p^window_corrected^* was used as the similarity score between the PSSMs.

### Sparsity

Sparsity measures the proportion of cells in the PSSM where there is no data. A sparsity of 1 denotes that all the cells in a PSSM have information and a sparsity of 0 denotes that there is no information in any of the cells in a PSSM.

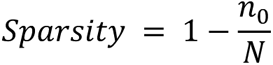

***Equation 2****: Sparsity ratio where* 𝑛_!_*is the number of cells in the PSSM with 0 value and N the total number of cells in the PSSM*.

### Establishing the optimal combination of selection days

All PSSMs split by the day of the selections (1,2,3 and 4) and combined day of the selections (1/2, 2/3, 3/4, 1/2/3 and 2/3/4) were compared with the dataset of 234 ELM class specificity determinant PSSMs. A similarity score *p*-value and the similarity score-derived rank of each comparison were calculated. The comparisons with the bait-peptide pair PSSM with the expected ELM class were classified as True Positive and all the other comparisons were classified as False Positives. For both the p-value and the rank data, a ROC analysis was performed and the area under the curve (AUC) was calculated to measure the quality of the selections on different days and combinations of days. The metrics were calculated using the *roc_curve* and *auc* functions from the *sklearn.metrics* library in Python 3.9.7 respectively.

### Replicate comparison

The specificity determinants derived from overlapping and distinct peptide replicates were compared for each peptide in the DMS-BM bait-peptide pair set and a comparison *p-value* was calculated from each comparison. The *p*-values for the comparison of the bait-peptide pairs were grouped based on the following criteria: (i) “Same bait / Overlapping peptide” for the bait-peptide pair with the same bait and overlapping peptides, (ii) “Same bait - Different peptide” for the bait - peptide pair with the same bait and non-overlapping peptides, and “Other” for the remaining peptides. “Same Bait - Same Peptide” were excluded from the analysis. The groups were then plotted as boxplots using the *seaborn 0.11.1* library in Python 3.9.7. The difference in the means of the groups was compared with a pairwise Mann-Whitney test using the *stats.mannwhitneyu* function from the *scipy 1.7.0* library in Python 3.9.7.

### Expression and purification of proteins for affinity measurements

His-GST-tagged human bait proteins were expressed in 4 L *E. coli* BL21 (DE3) cultures for fluorescence polarisation (FP) based affinity measurements. Lysate was cleared by centrifugation (16,000 RCF, 4°C, 1 h) and the supernatant was mixed with Ni Sepharose High Performance resin (Cytiva) (1 mL beads/pellet) followed by an incubation (4°C, agitation, 1 h). The beads were washed with washing buffer (20 mM NaPO_4_, 0.5 M NaCl, 30 mM imidazole, pH = 7.5). The His-GST-tags were cleaved by incubation for 16-18 h at 4 °C in 200 μL 0.5 mg HRV 3C protease and 1 mL primary buffer (20 mM NaPO_4_, 0.5 M NaCl, pH = 7.5). Cleaved proteins were collected. The samples were dialysed to 50 mM potassium phosphate buffer (pH = 7.5) for 16-18 h at 4°C. Protein purity and quality was confirmed through SDS-PAGE and thermal shift assay (Tycho NT.6, NanoTemper). His-GST tagged domains of SARS-CoV-2 proteins were expressed and harvested as described above. After the centrifugation step, the lysate was mixed with Pierce glutathione agarose (ThermoFisher) and incubated on 4°C under agitation for 30 minutes. The gel was washed with wash buffer 2 (50 mM Tris, 300 mM NaCl, 2 mM DTT, pH 7.8) and the protein of interest was eluted with elution buffer (wash buffer 2 supplemented with 10 mM reduced GSH). The His-GST tag was cleaved using HRV 3C protease (inhouse; 18 h at 4 °C) and the cleaved tag was removed by reverse immobilized metal affinity chromatography. Purified proteins were subjected to size exclusion chromatography (HiLoad 16/600 Superdex 75 pg; Cytiva) to remove any residual impurities, concentrated, flash frozen and stored at -80 °C until further use.

### FP-monitored affinity measurements

FP measurements were carried out in triplicates with an SpectraMax iD5 Multi-Mode Microplate Reader (Molecular Devices) in Corning 96-Well Half-Area Plates [Black, Flat-bottom, non-binding surface (Corning, Cat: 3993]), with excitation: 485 nm, emission and 535 nm in a total volume of 50 μL. Peptides were obtained at >95% purity (GeneCust). FITC-labelled peptides were dissolved in dimethyl sulfoxide (DMSO) and diluted 1:1000 in 50 mM KPO_4_ buffer (pH 7.5). Unlabeled peptides were dissolved in 50 mM KPO_4_ buffer (pH 7.5). Peptide concentrations were determined spectroscopically (FITC-labelled peptides: λ = 495 nm, unlabeled peptides: λ = 280 nm). For saturation experiments, bait proteins in 50 mM KPO_4_ pH 7.5 solution (For SARS-CoV-2 protein domains the assay buffer was supplemented with 0.05% Tween20 and 1 mM TCEP) were arrayed in serial dilution (Diluent: 50 mM KPO_4_, pH = 7.5; for SARS-CoV-2 proteins same adjustment of buffer was made as described above): 25 μL protein solution followed by addition of 25 μL peptide master mix (2 mM DTT and 10 nM labelled peptide in 50 mM KPO_4_ buffer, pH 7.5). For the displacement experiments, unlabeled peptides were arrayed in serial dilution: 25 μL unlabeled peptide solution followed by addition of 25 μL peptide master mix supplemented with the protein of interest at a concentration of 4x the K_D_ value. Data was analysed using GraphPad Prism (GraphPad Software, San Diego, California USA).

### Infections experiment

VeroE6 cells were infected with SARS-CoV-2 (MOI:0.5) for 1 hour at 37°C and 5% CO_2_, then inoculum was removed and replaced with medium containing the indicate concentration of peptide. Eight hours post infection cells were lysed and RNA was isolated using NucleoSpin RNA Plus XS (Macherey Nagel) according to the manufacturer’s instructions. cDNA was synthesized using High-capacity cDNA Reverse Transcription kit (Thermo Fisher). SARS-CoV-2 RNA was quantified using qPCRBIO probe mix Hi-ROX (PCR Biosystems) and the following primers and probes, GTCATGTGTGGCGGTTCACT, CAACACTATTAGCATAAGCAGTTGT and FAM-CAGGTGGAACCTCATCAGGAGATGC-BHQ. GAPDH was used as a reference gene, detected by RT qPCR Primer Assay (NM_001195426, Qiagen) and the qPCRBIO SyGreen mix Hi-ROX (PCR Biosystems). qPCR experiments were run on a StepOnePlus real-time PCR system (Applied Biosystems).

### Viability test

Cells were treated with the indicated peptides for 8 h, then cellular viability was determined using CellTiter-Glo® Luminescent Cell Viability Assay (Promega) on a Varioskan LUX Multimode Microplate Reader (ThermoFisher Scientific).

## Acknowledgments

We thank Jakob Nilsson for providing purified B56, and Elias Tjärnhage for providing the items for Fig. 1. This project was supported by the Swedish research council (YI/AÖ: 2022-05278, 2023; YI: 2020-03380; PJ:2020-04395) and Cancer Research UK (Senior Cancer Research Fellowship grant (C68484/A28159) to ND and IT). Sequencing was performed by the SNP&SEQ Technology Platform in Stockholm. The facility is part of the National Genomic Infrastructure (NGI) Sweden and Science for Life Laboratory and is also supported by the Swedish Research Council and the Knut and Alice Wallenberg. We thank Julia Vargas for insightful advices on AlphaFold modelling of protein-peptide interactions.

## Author contributions

CB, AKÖ, ND and YI conceived and designed the research project. LM and CB performed phage display and library generation affinity measurements and initial data analysis. FM performed affinity measurements under supervision of PJ. RL performed the *in vitro* assay. LS, IT, and ND performed data analysis. LS generated all final figures. CB and YI wrote the manuscript with the input of all other authors.

## Supplemental information

**Figure S1.**
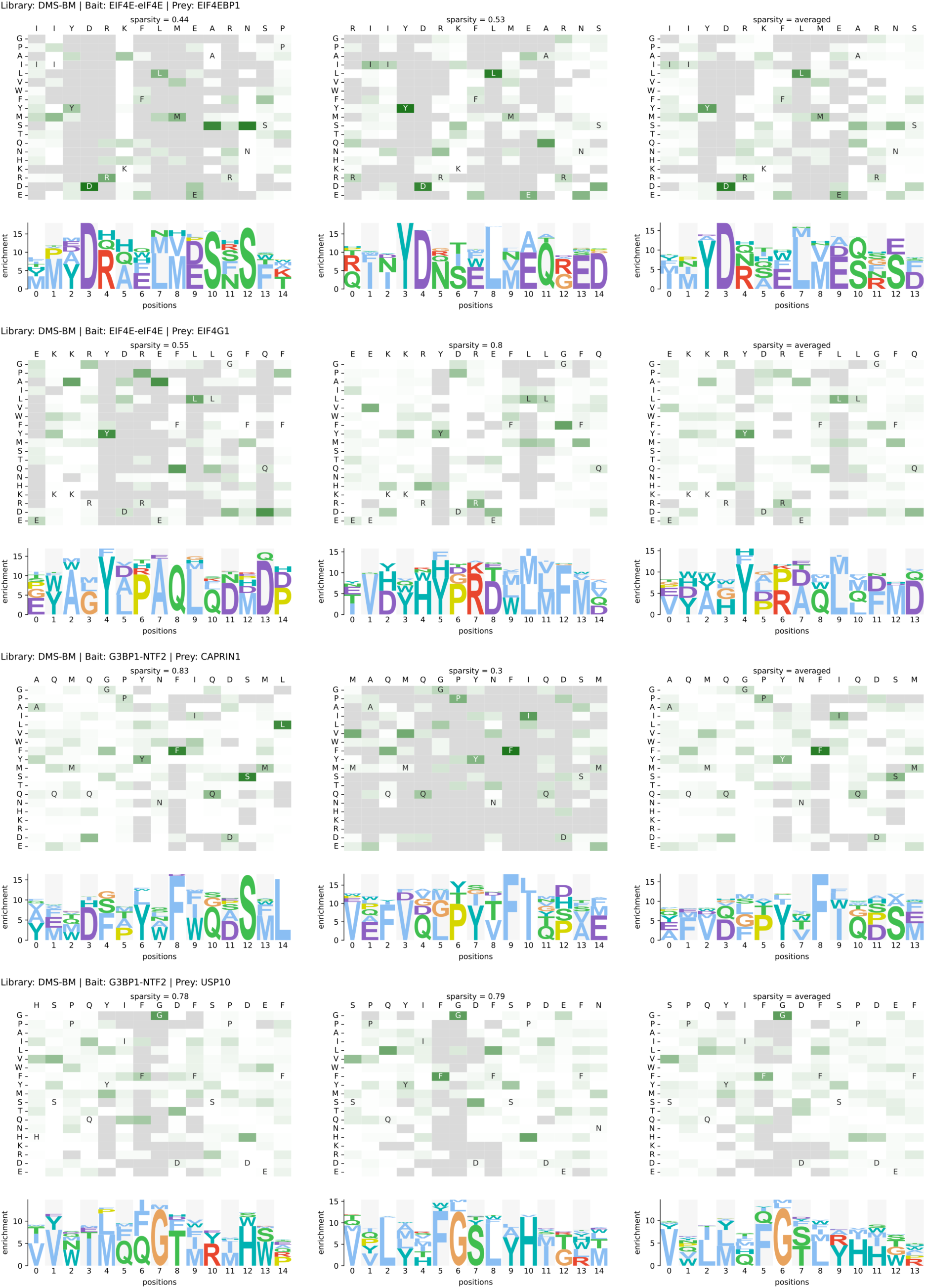
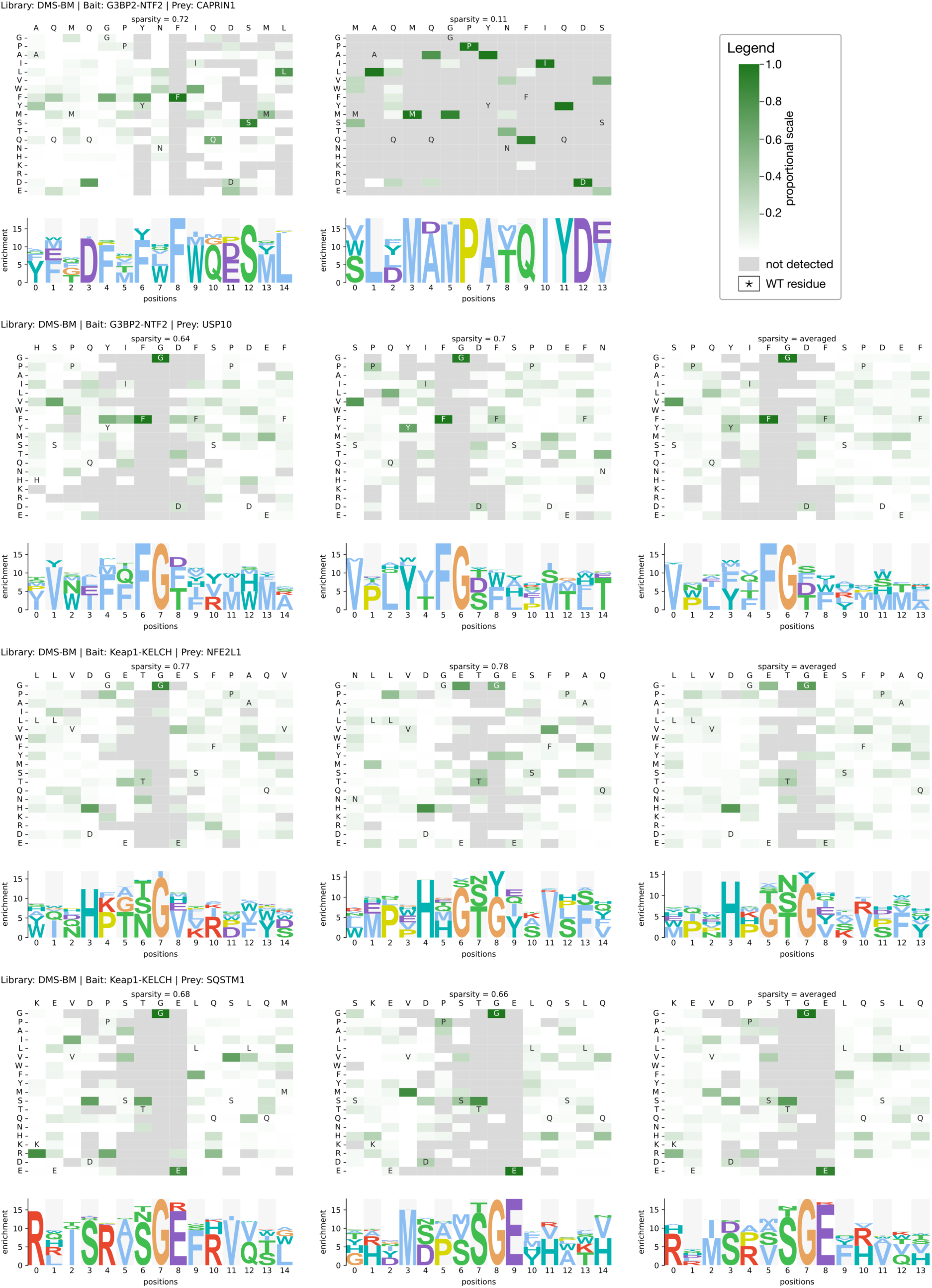
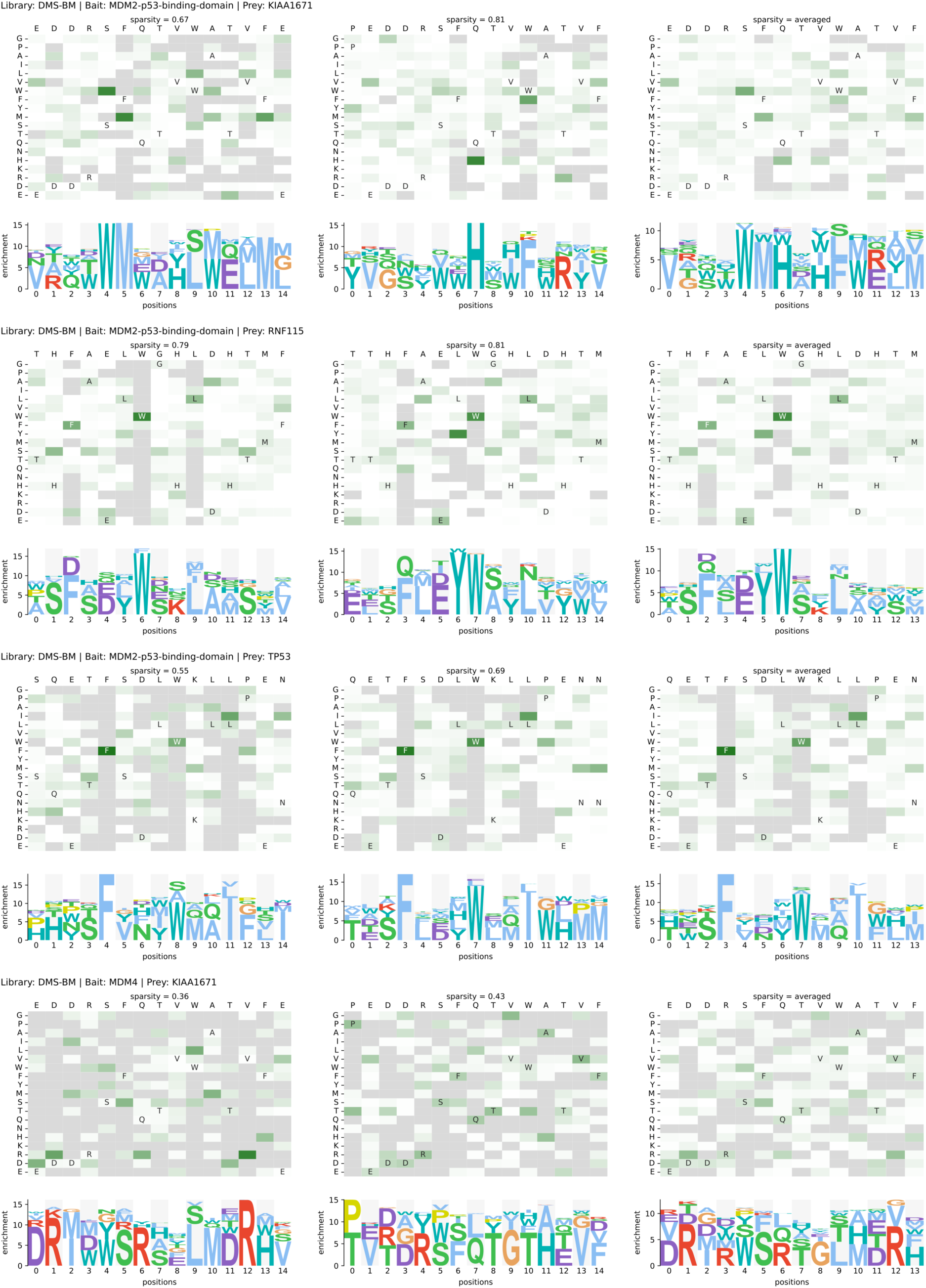
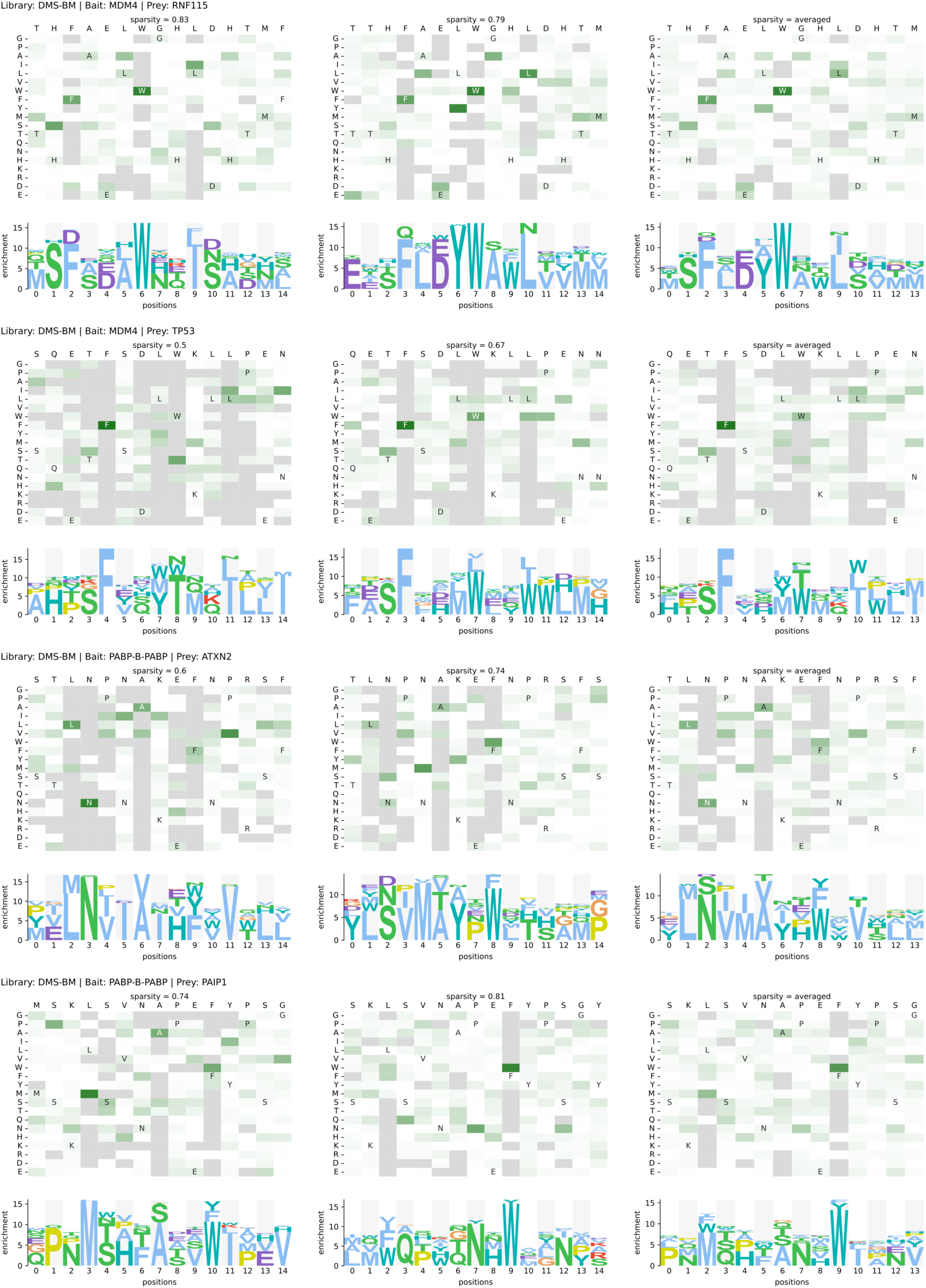
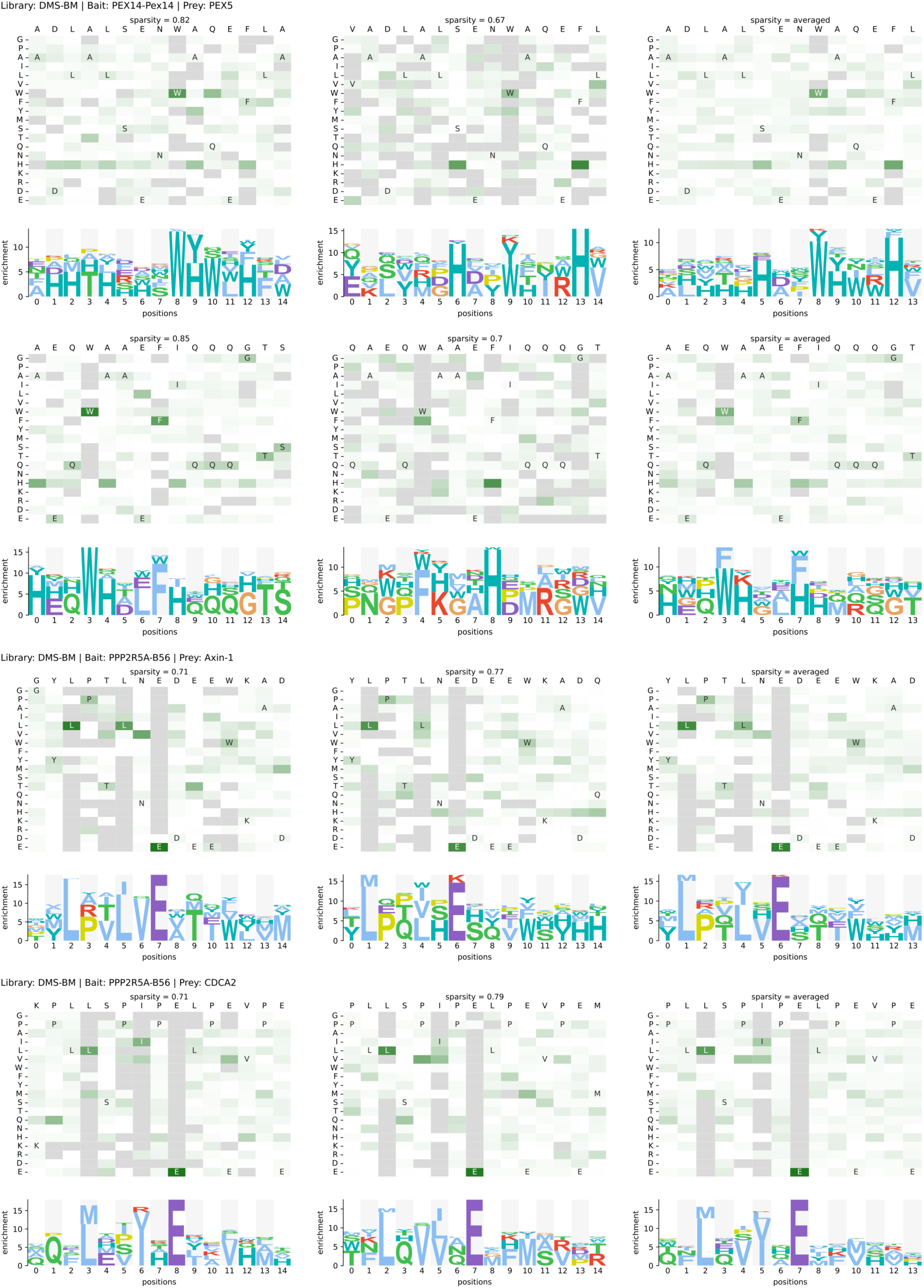
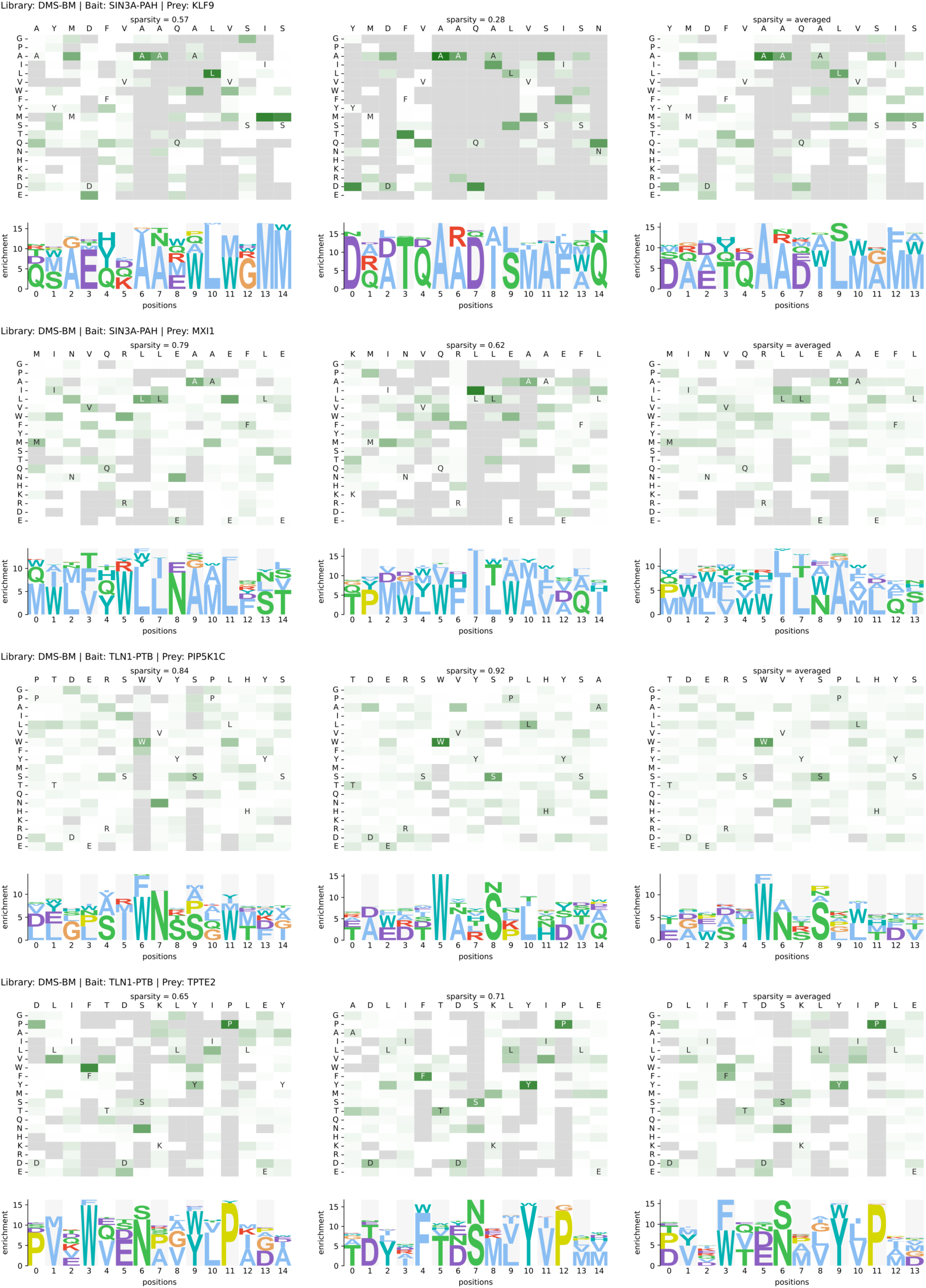
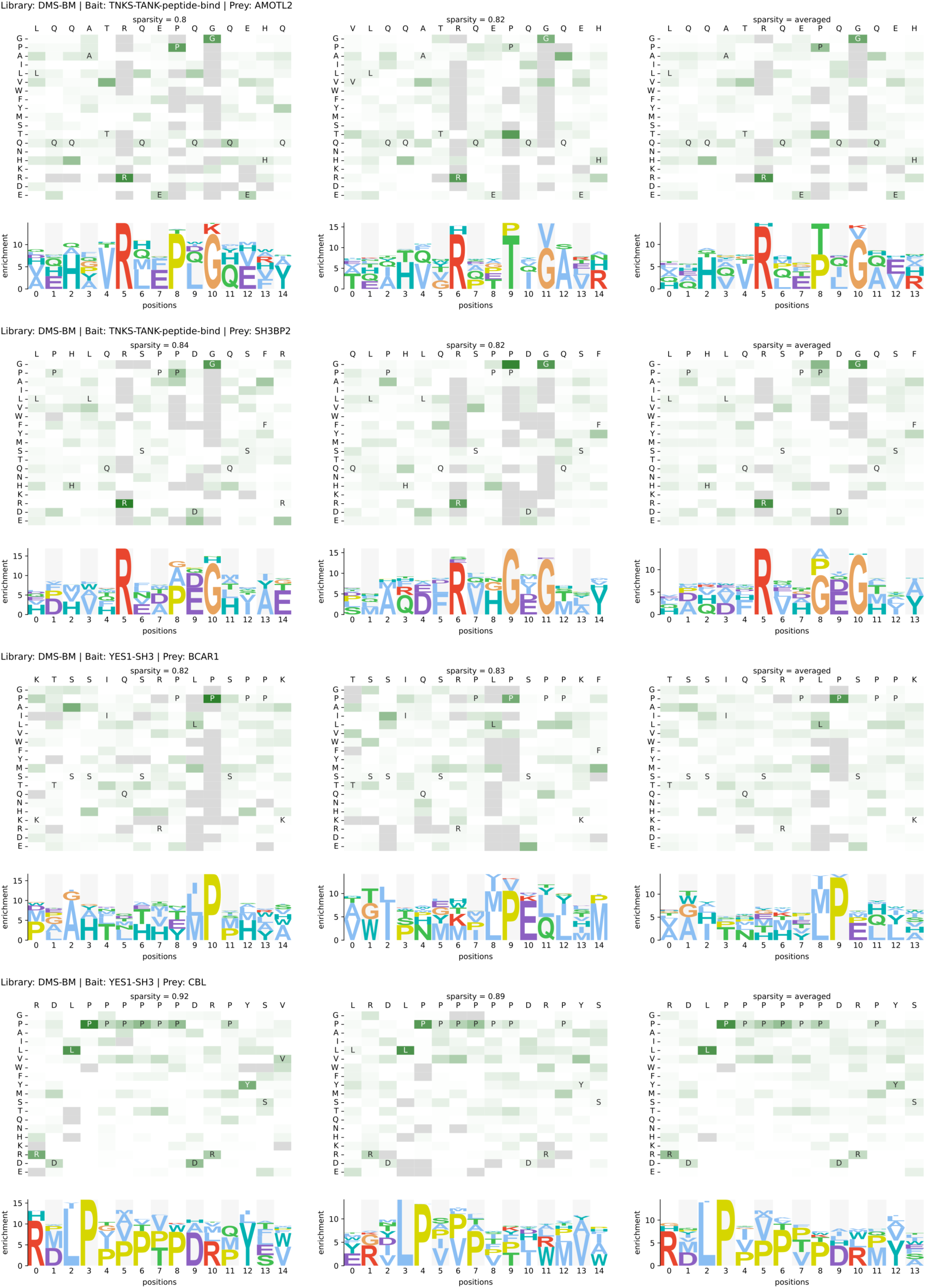
Heat map and PSSM representation of DMS analysis results generated through selections against the DMS-BM library. Each parental peptide is represented twice (left and middle) in overlapping registry. The averaged results are showed to the right.

**Figure S2.**
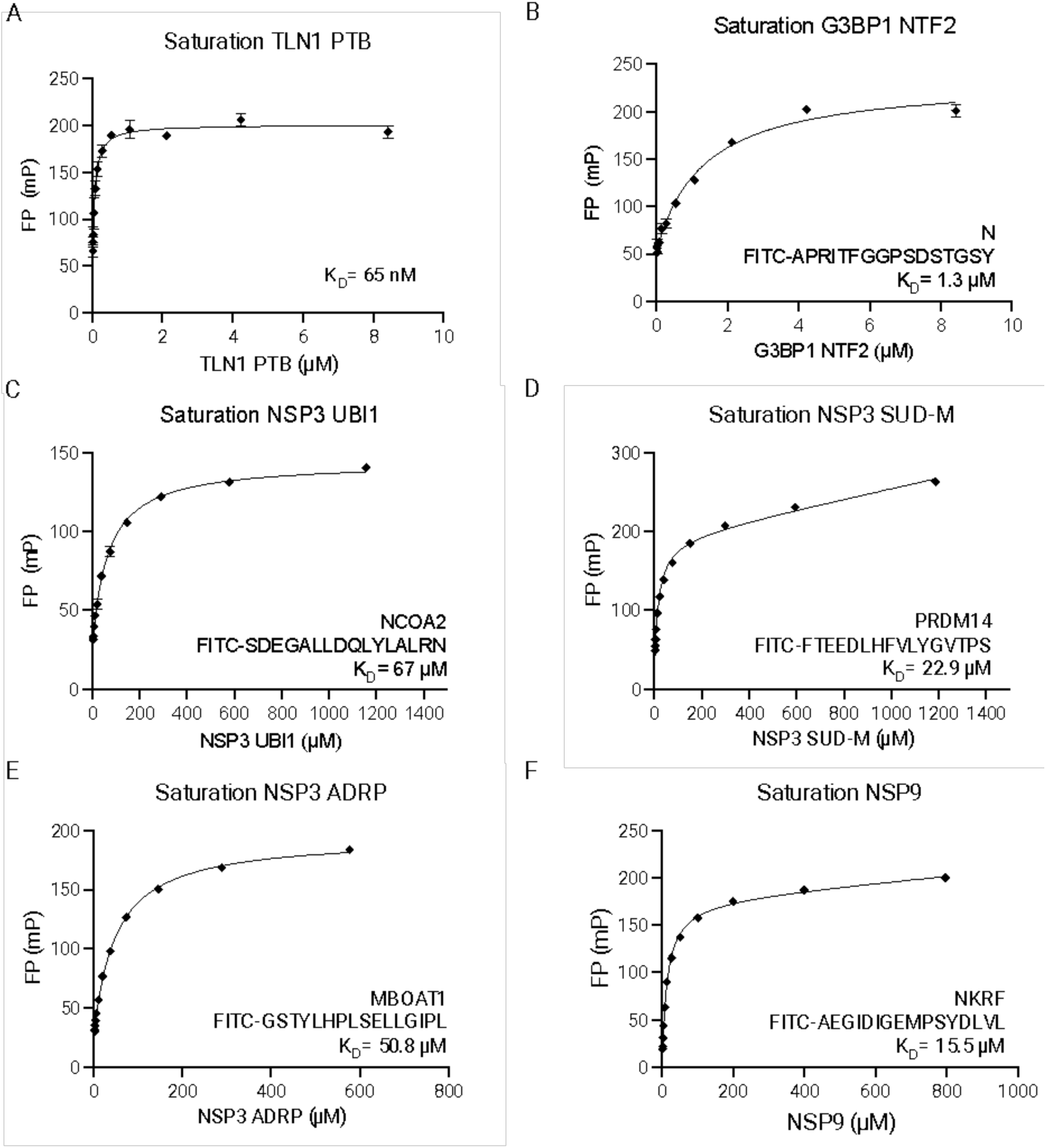
Binding titration curves as detected by FP of FITC-labeled probe peptides (indicated) binding to A) TLN1 PTB, B) G3BP1 NTF2, C) NSP3 UBl1, D) NSP3 SUD-M, E) NSP3 ADRP, and F) NSP9.

**Figure S3.**
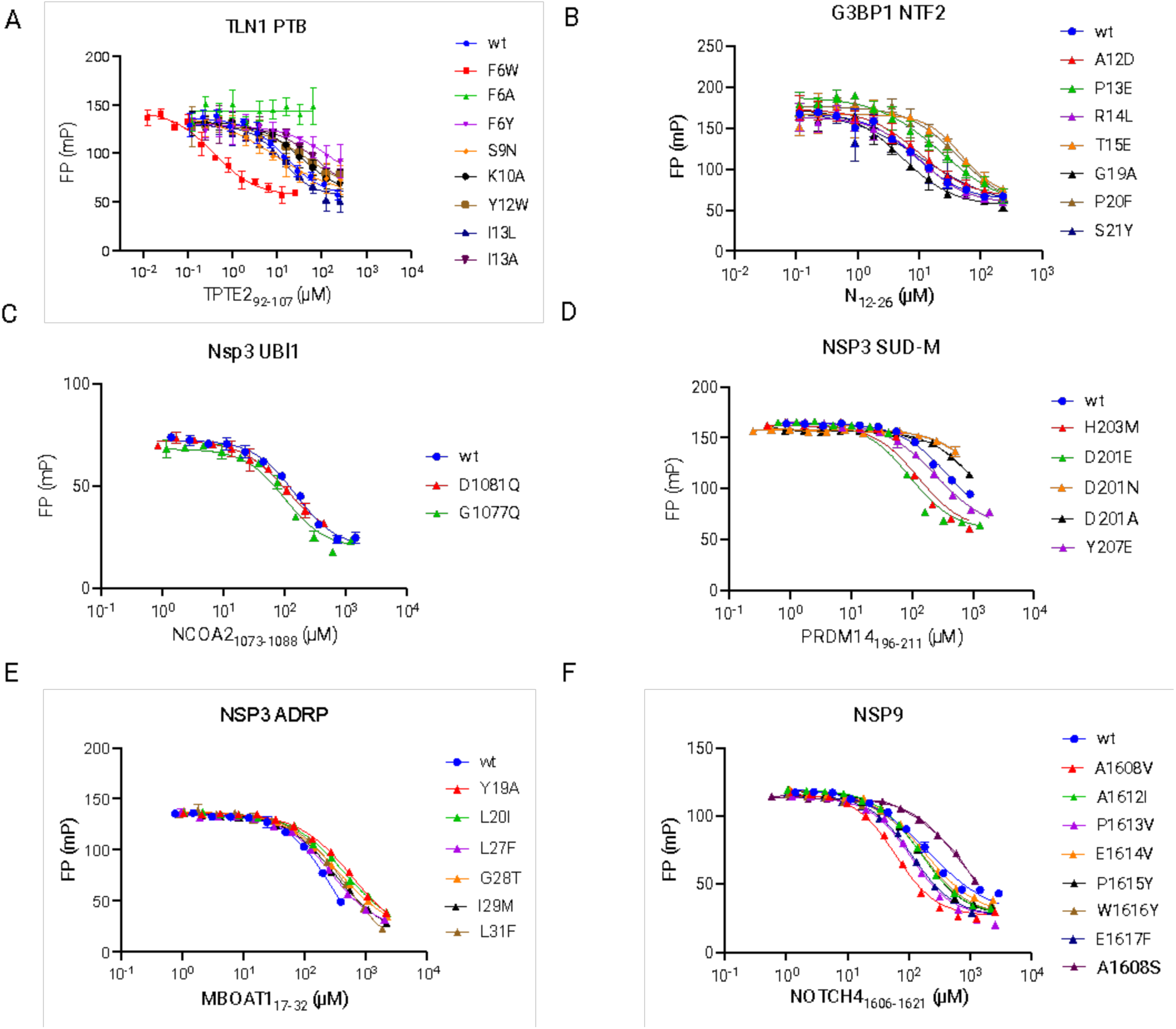
Affinity determinations through fluorescence polarization-based competition experiments of wild-type and mutant TLN1 PTB, B) G3BP1 NTF2 ,C) NSP3 UBl1, D) NSP3 SUD-M, E) NSP3 ADRP, and F) NSP9.

**Figure S4.**
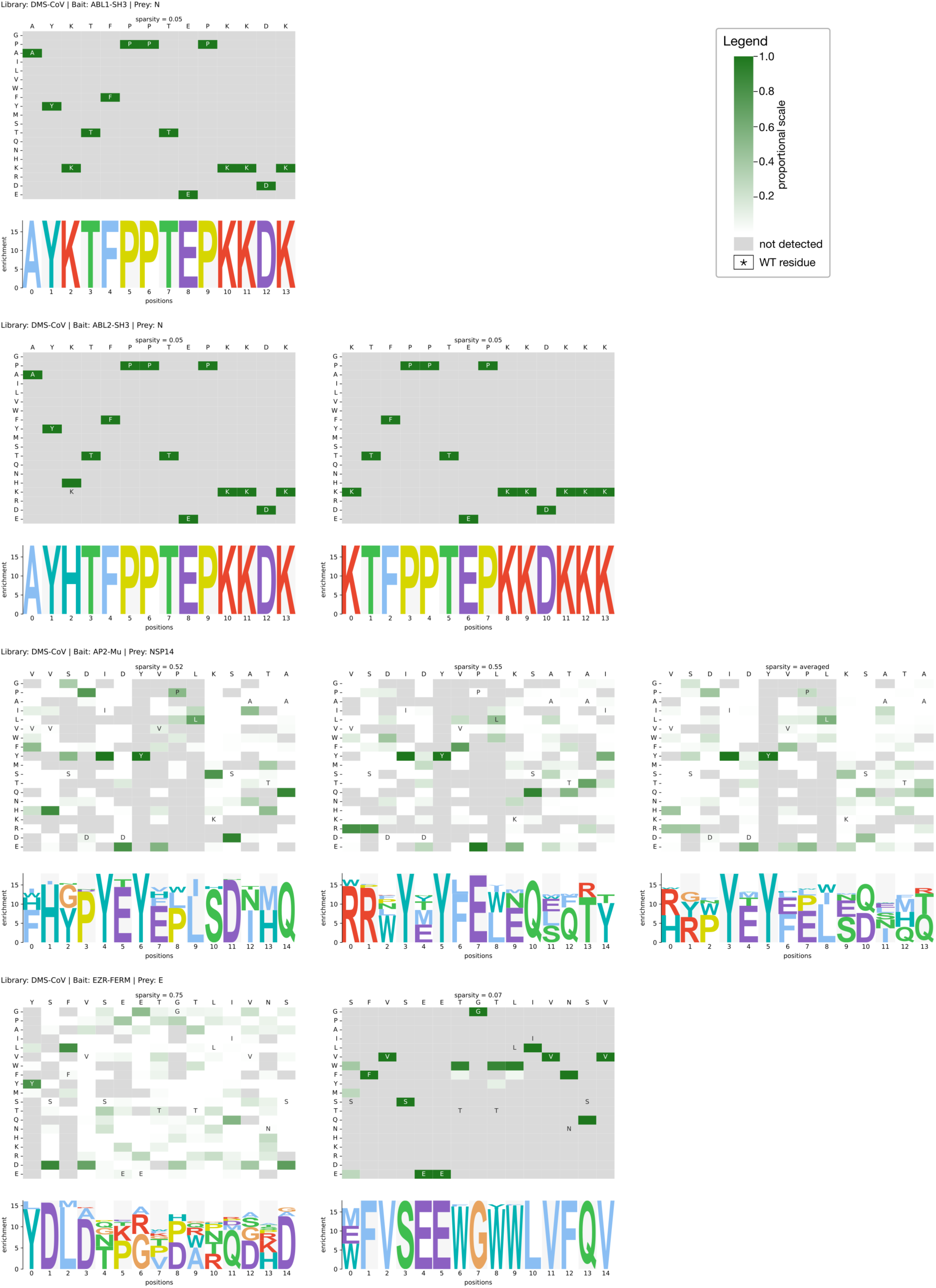
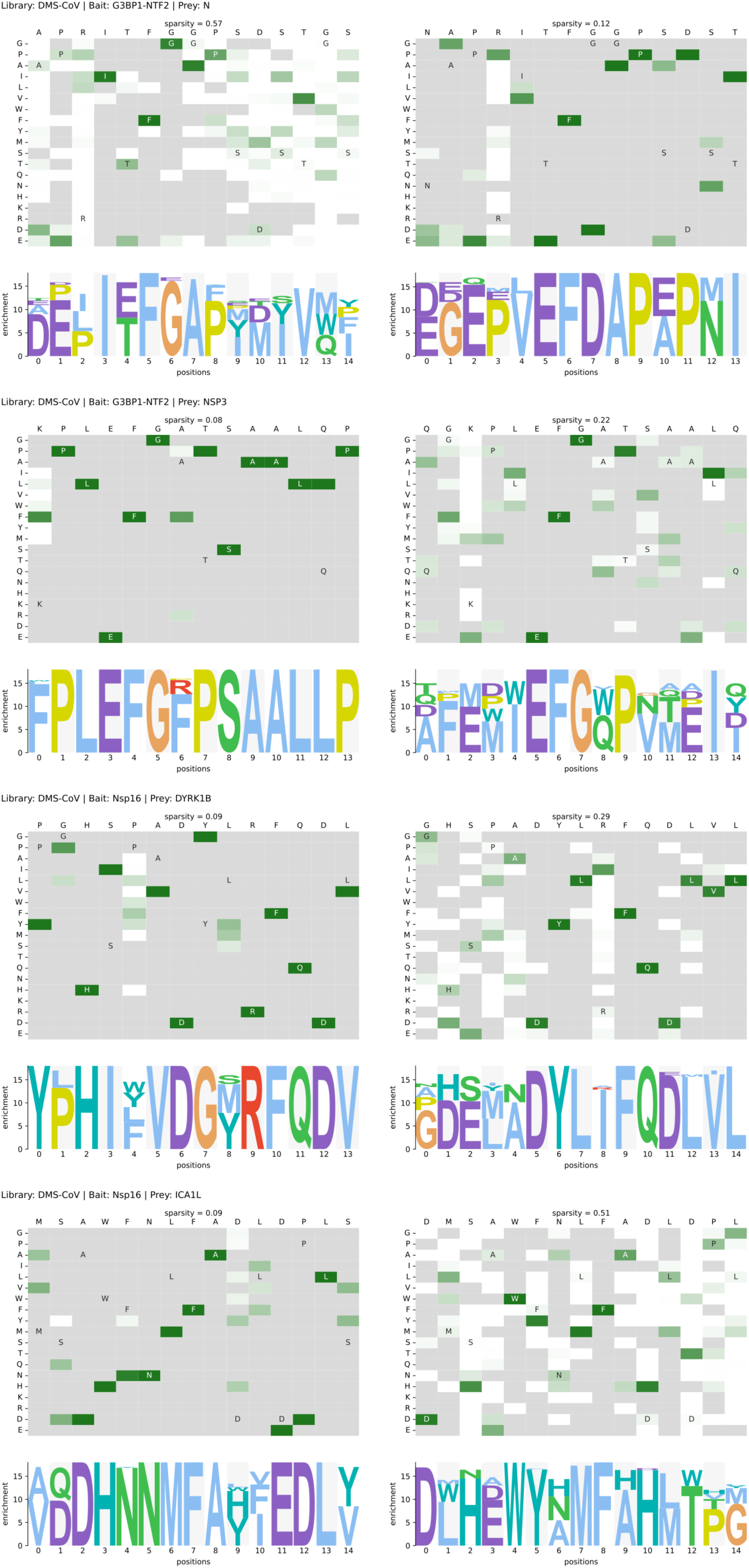
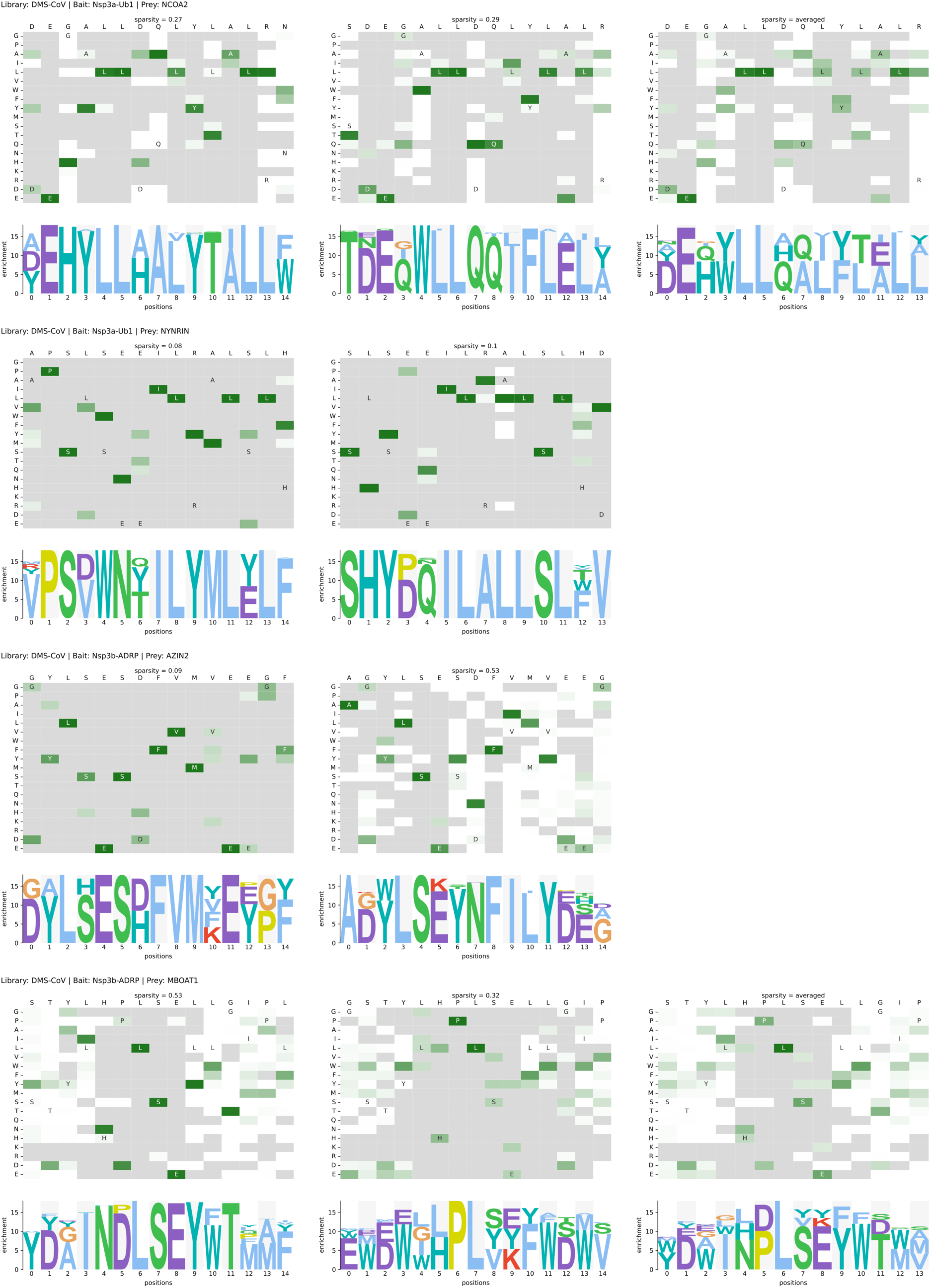
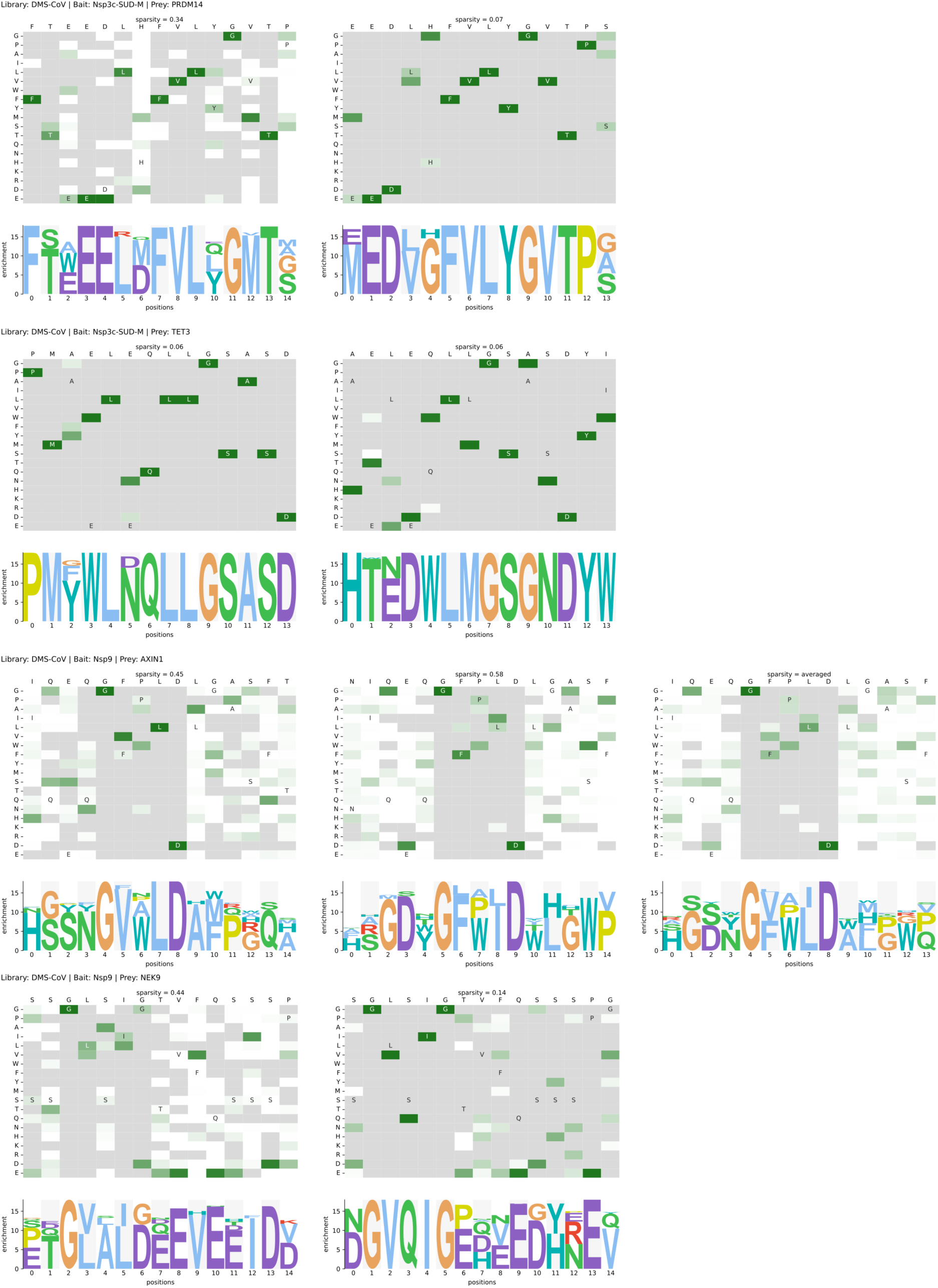
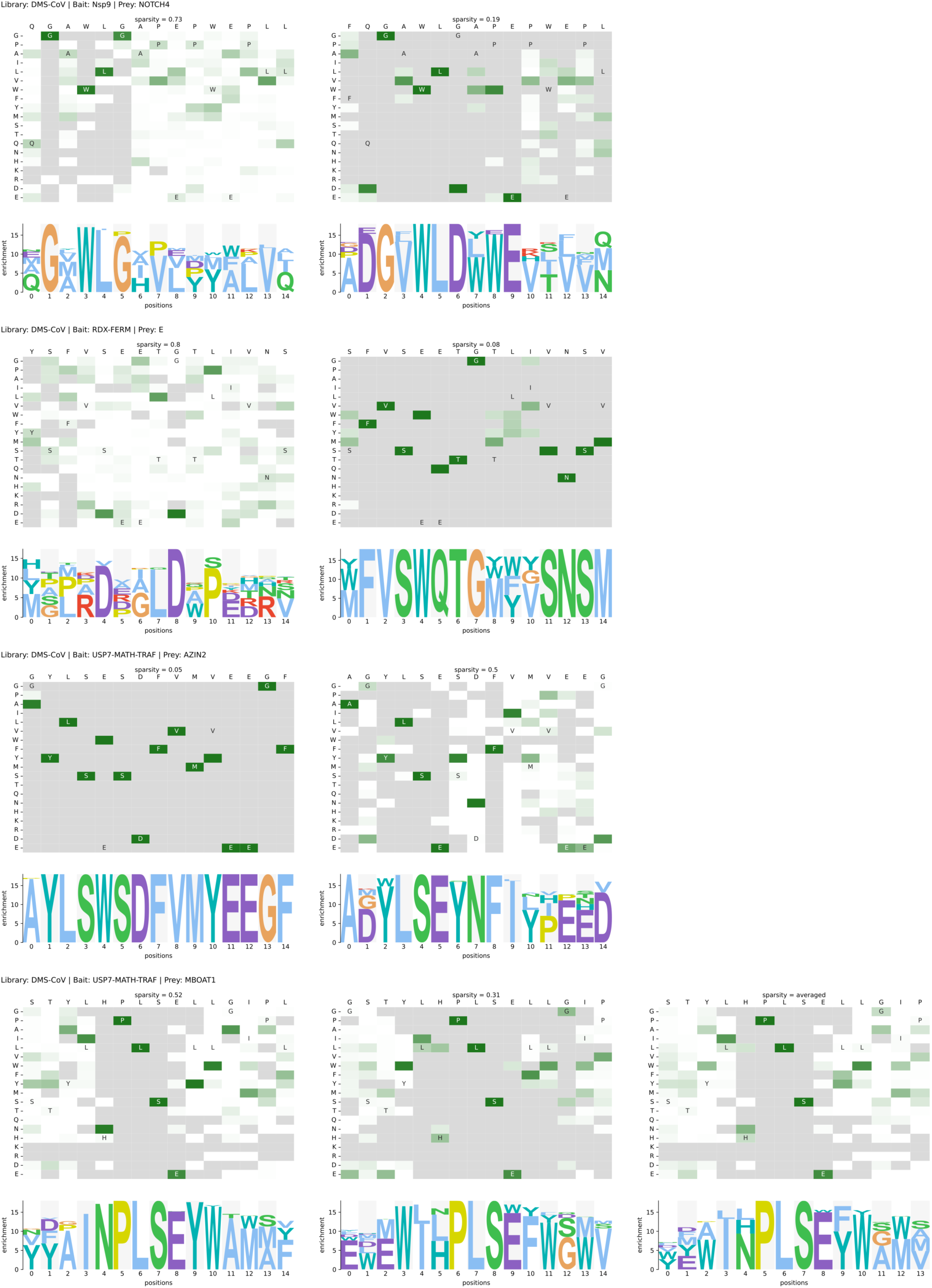
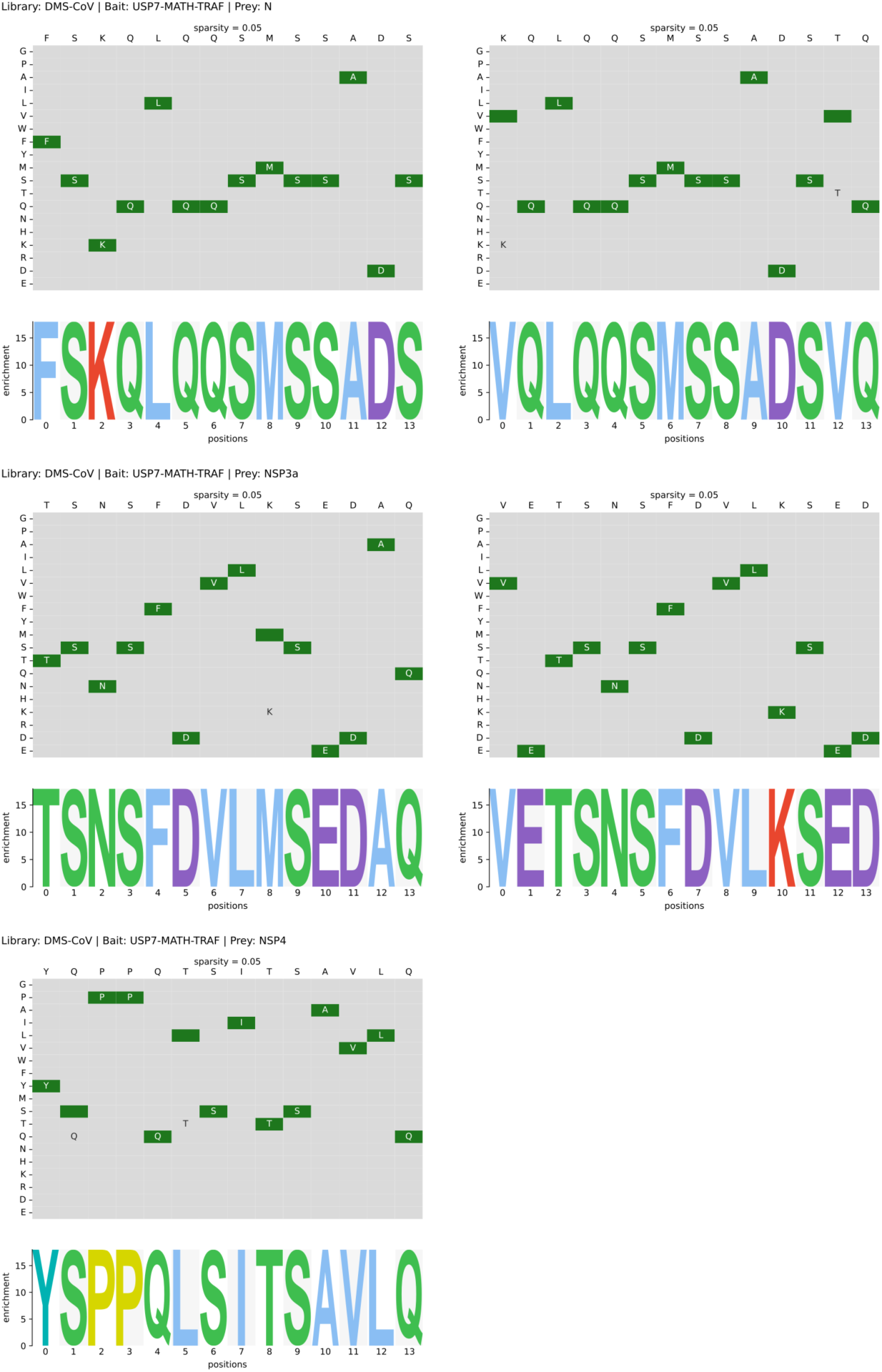
Heat map and PSSM representation of DMS analysis results generated through selections against the DMS-CoV library. Each parental peptide is represented twice (left and middle) in overlapping registry. The averaged results are showed to the right when available.

**Figure S5.**
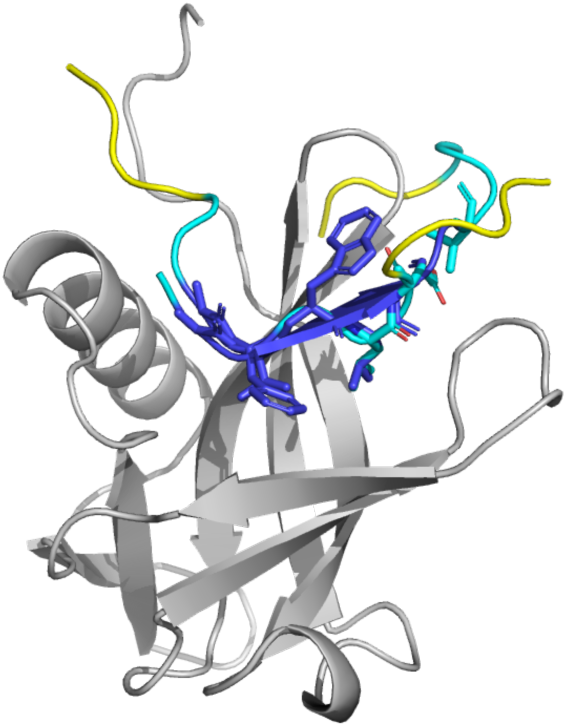
AlphaFold3 model of the complex of NSP9 and the NOTCH4 A1603V/P1613V peptide (QGVWLGAVEPWEPL) overlayed with the model of the AXIN (IQEQGFGFPLDLGAS). Peptide coloring is according to the pLLDT score (deep blue = high confidence). Generated using PyMol.

